# Oxidations and amino acid substitutions in urinary proteins are the distinguishing characteristics of aging

**DOI:** 10.1101/2020.07.13.199984

**Authors:** Yongtao Liu, Xuanzhen Pan, Yuanrui Hua, Yunlong Wang, Youhe Gao

**Affiliations:** Department of Biochemistry and Molecular Biology, Beijing Key Laboratory of Gene Engineering Drug and Biotechnology, Beijing Normal University, Beijing, China

**Keywords:** urine proteome, aging, global chemical modification, oxidation, amino acid substitution

## Abstract

Aging is an inevitable course of life. Additionally, the risk of chronic diseases or cancer increases with age. The comprehensive identification of signs related to aging can be beneficial for the prevention and early diagnosis of geriatric diseases. The comparison of global modifications in the urine proteome is a means of multidimensional information mining. This approach is based on urine, in which changes from whole-body metabolism can accumulate. This study used the urine of healthy people at different ages (22 children, 10 young people, 6 senior people) as the research object and using high-resolution tandem mass spectrometry, label-free quantitation combined with non-limiting modification identification algorithms and random group test, compared the differences in protein chemical modifications among three groups. The results show that multi-sites oxidative modifications and amino acid substitutions are noticeable features that distinguish these three age groups of people. The proportion of multi-site oxidations in urine proteins of senior (29.76%) is significantly higher than the young group (13.71% and 12.97%), which affect the biological processes of various proteins. This study could provide a reference for studies of aging mechanisms and biomarkers of age-related disease.

## 0. Introduction

Owing to various biochemical reactions of proteins, such as synthesis, catalysis, and regulation, biological processes can operate simultaneously and coordinately in living organisms. Proteins are biological macromolecules with complex structures, and differences in their advanced structures lead to different biochemical activities. As a result, proteins combine into networks to perform specific functions [1]. The chemical modification of a protein refers to reactions of amino acid residues or the covalent group at the end of the chain, and modification usually changes the advanced structure and function of the protein. A few of these chemical structure changes do not affect the biological activity of the protein, which are called the modification of non-essential parts of protein, but in most cases, the change of molecular structure will significantly change the physical and chemical properties of the protein, bring great changes to the conformation and activity of the protein, and thereby influence the function it performs [2]. Therefore, even if the protein content level does not change, if there are some small changes in the chemical modification level, the protein function will significantly change, thus, the chemical modification enriches the functions and regulations of the protein in another dimension. It is often observed that inactive and abnormal proteins accumulate in older cells and tissue [3–5]. The effects of chemical modification on protein function are mainly reflected in the following three aspects: 1. Even if a protein site is modified, its function will be affected. 2. For the same protein, the same or different modifications of different amino acids affect their functions differently. 3. The same protein may undergo multiple types of chemical modifications, which in turn help it to participate in biological processes in a more varied and complex way [6]. The main types of chemical modifications related to proteins are as follows: 1. Post-translational modification (PTM) [7], refers to the chemical modification of a protein after its translation. The precursor protein prior to post-translational modification usually has no biological activity, and needs to be modified after translation by specific modifying enzymes before it can become a functional mature protein and exert its specific biological functions [8, 9]. 2. Chemical derivative modification of a protein refers to modification by the introduction of new groups into the side chain or the removal of existing protein groups, which usually includes spontaneous non-enzymatic modifications and modifications introduced by cross-linking agents and artificial reagents. 3. Amino acid substitution refers to changes in protein properties and functions caused by the original amino acids in protein side chains being replaced by other amino acids. These modification types can significantly affect protein functions.

Mass spectrometry can not only achieve large-scale omics data acquisition and deep mining but also accurately determine the targeted modification of specific proteins. With the continuous development of instrument science and technology, ultrahigh-resolution tandem mass spectrometry (LC-MS/MS) provides richer and more accurate information and data for proteomics and modification research and also provides powerful help for accurately identifying chemical modification sites in protein chains. Generally, when analysing proteomic data, it is necessary to study and compare species proteome databases. When using search engines to search databases, it is usually necessary to manually set known types of protein chemical modification. This type of search is called the limiting search. However, when the types of protein modification in a sample are unknown or when researches are hoping to find a new type of modification, limiting search is restricted [10]. Therefore, a comprehensive, non-limiting modification search is crucial to understanding all the chemical modification information contained in the proteome of a sample. The Open-pFind algorithm is an open sequence library search algorithm that integrates information from the UniMod database, using an open search method to analyse and process the collected mass spectrometry data [11–13]. This is an innovative idea to obtain the global chemical modification information of the proteome.

Although the total amount of protein in healthy people’s urine is relatively low, it is extremely rich in species. It is known that more than 6,000 different proteins can be identified in normal human urine [14]. Under normal circumstances, glomerular proteins can be screened from the initial urine, 98% is reabsorbed in the renal tubules, and the remaining 2% protein is excreted together with a small amount of mucin secreted by the renal tubules and other urothelial cells. Urine is generated in the kidney and stored in the bladder for several hours. Therefore, the protein information in the urine can more directly reflect the urinary system. However, most of the proteins are collected in urine after the glomerular filtration of plasma and reabsorption in renal tubules. In theory, these proteins all come from blood. They can still be retained after kidney filtration and reabsorption, which is enough to show that the remaining 2% protein is not randomly or non-selectively “leaked” from a healthy kidney. Chemical modification is another important research direction of urine proteomics. Overall comparison in the chemical modification level of the urine proteome will also present more abundant information for physiological changes in living organisms [9]. At present, there are more than 1,500 chemical modifications of protein in UniMod [15], PSI-MOD, and RESID databases. Plasma proteome can reflect the subtle situation between age and senility [16]. As another body fluid rich in protein information-the urine, it has been found that urine proteome can also reflect the aging information of the human body [17]. In our previous study on the plasma proteome of newborn rats [18], we also found significant differences in different developmental stages. Even clues about common physiological processes such as hunger can be found in the urine proteome [19]. Based on this and the importance of chemical modification in the urine proteome of different age groups, the purpose of this study is to compare the global chemical modification levels of the urinary proteome of three groups by high-resolution tandem mass spectrometry combined with non-limiting modification search (Open-pFind).

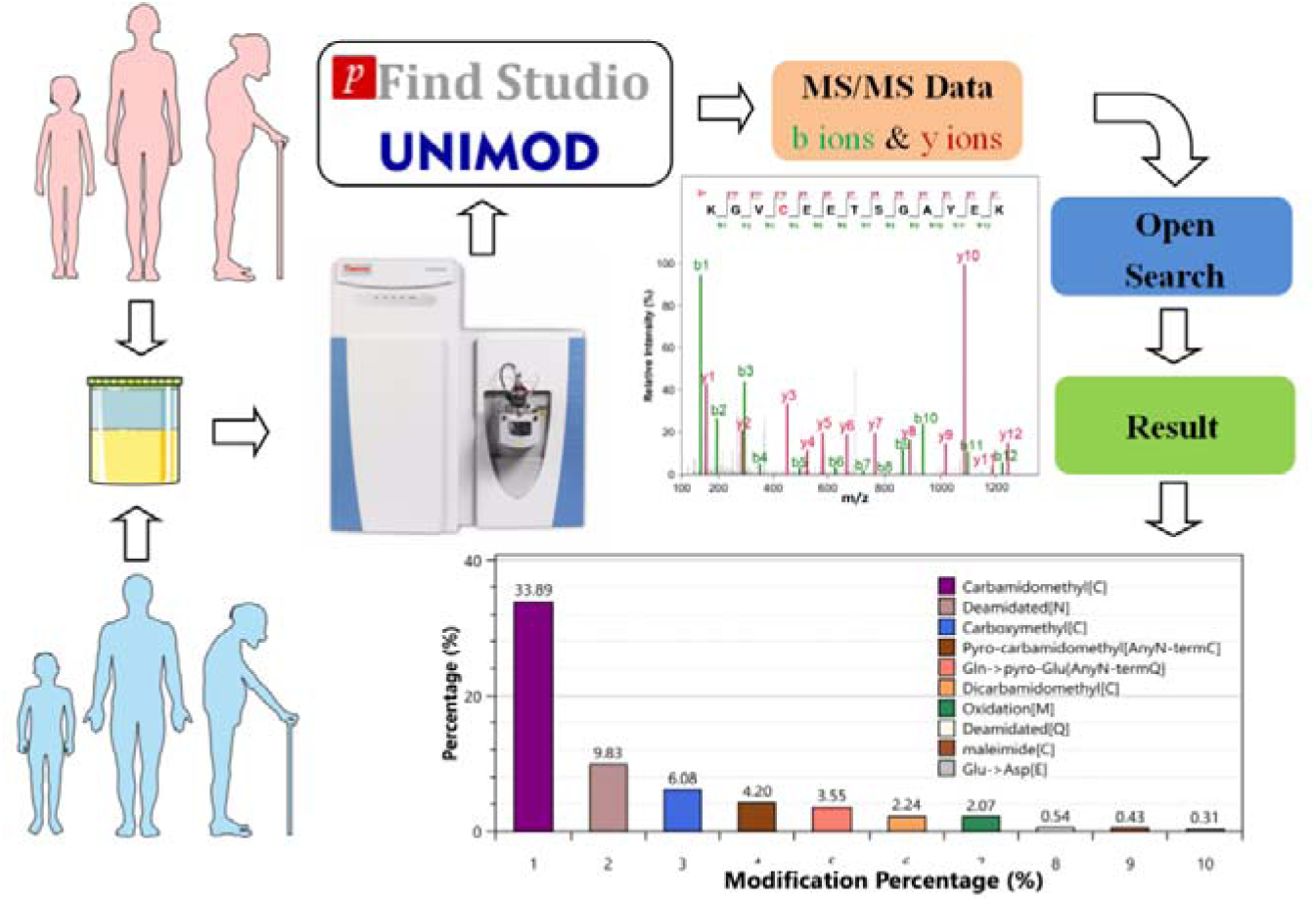

## 1. Materials and methods

### 1.1 Samples and data sources

22 cases of children urine proteome data came from a study on patients with vesicoureteral reflux (VUR) (original data download address: http://proteomecentral.proteomexchange.org/cgi/GetDataset?ID=PXD010469). 6 cases of senior urine proteome data came from a study of neutrophil-associated inflammation in the urinary tract (Raw data download address http://proteomecentral.proteomexchange.org/cgi/GetDataset? ID=PXD004713). The data downloaded above are all healthy control samples. 10 cases of healthy young age urine proteome data came from previous articles published in our laboratory. The samples are taken from our laboratory volunteers, and there were no restrictions or requirements on the volunteers’ diet, drugs, and other factors. The samples’ information is shown in **Table 1**. Raw data of mass spectrometry and results of pFind were submitted to iProX Datasets (https://www.iprox.org/page/HMV006.html) under the Project ID: IPX0002313003.

**Table 1.** Information and statistics of the downloaded raw data

|  | Healthy urine sample (n=38) |  |  |
| --- | --- | --- | --- |
|  | Children (n=22) | Young age (n=10) | Senior (n=6) |
| Gender |  |  |  |
| Male | 13 | 5 | 2 |
| Female | 9 | 5 | 4 |
| Age |  |  |  |
| Average ± standard deviation | 5.7±3 | 26.4±2 | 76.5±5 |
| Age range | 1~22 | 22~29 | 70~81 |

### 1.2 Protein sample preparation and trypsin digestion

This section only introduces the young age group sample processing methods in detail. The urine samples were reacted with 20 mmol/L dithiothreitol (DTT) at 37 □ for 1 h to denature the disulfide bonds in the protein structure, followed by the addition of 55 mmol/L iodoacetamide (IAA) in the dark for 30 min to alkylate the disulfide bond site. Precipitate the supernatant with three-Fold volumes of pre-cooled acetone at −20°C for 2∼4h, and then centrifuge at 12,000×g for 30 min at 4°C to obtain protein precipitate. The pellet was then resuspended in an appropriate amount of protein solubilization solution (8 mol/L urea, 2 mol/L thiourea, 25 mmol/L DTT, and 50 mmol/L Tris). The protein concentrated solution was measured using Bradford analysis. By using filter-assisted sample preparation (FASP) method [20], 100 µg of each sample was digested with trypsin (Trypsin Gold, Mass Spec Grade, Promega, Fitchburg, WI, USA) at a ratio of 50:1. After enzymolysis at 37°C for 14 h, 10% formic acid solution was added to the solution to terminate the enzymolysis, and the peptide solution was obtained after centrifugation through a 10 KDa ultrafiltration tube. The concentration of the peptide was determined using the BCA method and passed through a vacuum centrifugal concentrator (Thermo Fisher, USA), and the dried peptides are sealed and stored at −80°C. **Table 2** shows the urine sample processing methods of the other two groups in the literature, and the comparison with this method.

**Table 2.** Comparison of treatment methods of urine samples

|  | Children's urine | Young age's urine | Senior's urine |
| --- | --- | --- | --- |
| Sample pretreatment | 4500 × g, centrifuged for 20 min; | 2500 × g at 18 °C, centrifuged for 15 min; | 2500 × g at 10 °C, centrifuged for 15 min; |
| Protein enrichment and concentration | Centrifugal enrichment with 10 kDa filter; | Pre-cooled acetone precipitation enrichment; | Centrifugal enrichment with 10 kDa filter; |
| Reduction and alkylation methods | DTT/IAA; | DTT/IAA; | DTT/IAA; |
| Enzymatic hydrolysis of protein | Solid-phase Reversible Sample-preparation (SRS); | Filter-assisted sample preparation (FASP); | Filter-assisted sample preparation (FASP); |
| Type of protease | Trypsin; | Trypsin; | Trypsin; |
| Peptide desalting | Not mentioned; | C18 SPE ; | Stage tip ; |

### 1.3 Liquid chromatography-tandem mass spectrometry (LC-MS/MS) analysis and database search

Before analysis of urine samples of healthy young age, the dried peptide samples should be dissolved in 0.1% formic acid, the final concentration should be controlled at 0.1 μg/μL, and each sample should be analyzed with 1 μg peptide: Thermo EASY-nLC1200 chromatography system is loaded to Pre-column and the analytical column. Proteome data was collected by the Thermo Orbitrap Fusion Lumos mass spectrometry system (Thermo Fisher Scientific, Bremen, Germany). Liquid chromatography analysis method: pre-column: 75 μm×2 cm, nanoViper C18, 2 μm, 100Å; analytical column: 50 μm×15 cm, nanoViper C18, 2 μm, 100 Å; injection volume: 10 μL, flow rate: 250 nL/min. The mobile phase configuration is as follows, phase A: 100% mass spectrometric grade water (Fisher Scientific, Spain)/1‰ formic acid (Fisher Scientific), phase B: 80% acetonitrile (Fisher Scientific, USA)/20% water/1‰ formic acid, 120 Min gradient elution: 0 min, 3% phase B; 0 min-3 min, 8% phase B; 3 min-93 min, 22% phase B; 93 min-113 min, 35% phase B; 113 min-120 min, 90% phase B; mass spectrometry method, ion source: nanoESI, spray voltage: 2.0 kV, capillary temperature: 320C, S-lens RF Level: 30, resolution setting: level 1 (Orbitrap) 120,000 @m/z 200, Level 2 30,000 (Orbitrap) @m/z 200, precursor ion scan range: m/z 350-1350; product ion scan range: from m/z 110, MS1 AGC: 4e5, charge range: 2-7, Ion implantation time: 50 ms, MS2 AGC: 1e5, ion implantation time: 50ms, ion screening window: 2.0 m/z, fragmentation mode: high energy collision dissociation (HCD), energy: NCE 32, Data-dependent MS/MS : Top 20, dynamic exclusion time: 15s, internal calibration mass: 445.12003. **Table 3** shows the urine sample data collection methods of the other two populations reported in the literature and compares them with our method.

**Table 3.**
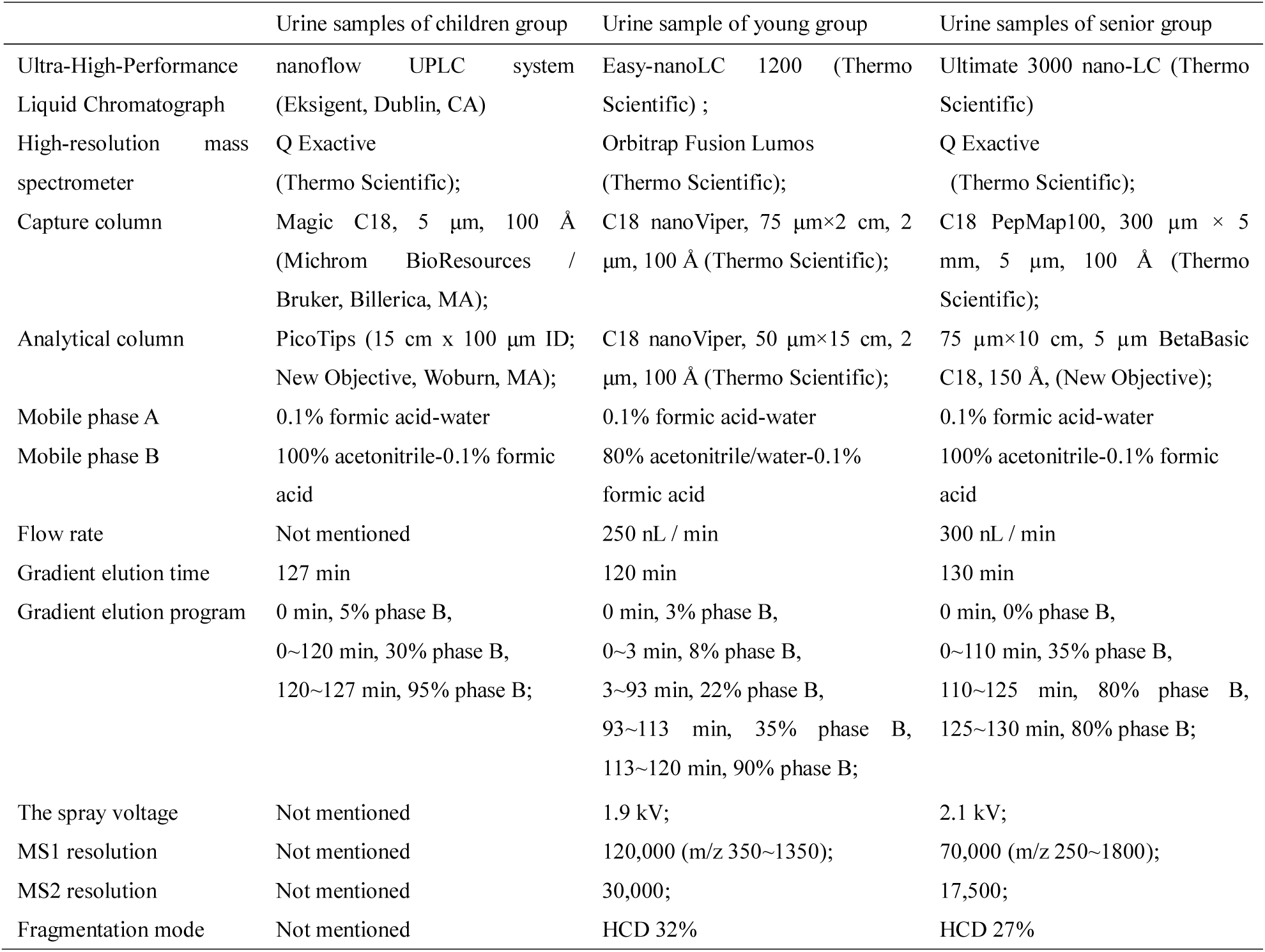
Comparison of instrument analysis methods for urine proteome samples

Use pFind Studio software (Version 3.1.6, Institute of Computing Technology, Chinese Academy of Sciences) to carry out a label-free quantitative analysis of LC-MS/MS data. The target search database is from the Homo Sapiens database (updated to February 2020) in Uniprot. Raw files generated by the Orbitrap Fusion were searched directly using a 20-ppm precursor mass tolerance and a 20-ppm fragment mass tolerance. While searching, the instrument type is HCD-FTMS, the enzyme is fully specific trypsin, up to 2 missed cleavage sites are allowed, and “open-search” is selected. Screening conditions: the FDRs were estimated by the program from the number and quality of spectral matches to the decoy database; for all data sets, the FDRs at spectrum, peptide, and protein level were <□1%, and Q value at the protein level is less than 1%. Data is analyzed using both forward and reverse database retrieval strategies.

### 1.4 Statistics and Analysis

Perform descriptive statistical calculations and analysis on the data provided by pFind, such as the median (for skewed distribution data) and proportion (percentage) of the quartile range. T-test, analysis of variance, Mann-Whitney U test, and Kruskal-Wallis test were used to evaluate the statistical comparison between children group, young group, and senior group. Statistical analysis was performed using GraphPad Prism v7.04. A p-value <0.05 is considered as significant.

## 2. Result

### 2.1 Identification of total protein by using bottom-up proteomics technology

Relying on the method of the label-free quantitative proteome, the experimental results of 38 samples were obtained by LC-MS/MS analysis. After retrieving data (raw) based on open-pFind software, the analysis results are shown in pBuild. The results are sorted and counted. The identification results of proteins and peptides in the samples are shown in **Table 4**.

**Table 4.** The results of proteins and peptides identification in urine samples

|  | Spectra | Scans | Peptides | Sequences | Protein Groups |
| --- | --- | --- | --- | --- | --- |
| Children Group | 29239 ± 5406 | 24763 ± 3833 | 13003 ± 2892 | 10002 ± 2252 | 2787 ± 898 |
| Young Group | 57001 ± 4820 | 42944 ± 2670 | 18024 ± 1562 | 12743 ± 1376 | 3447 ± 811 |
| Senior Group | 27007 ± 6575 | 23651 ± 5065 | 7906 ± 1899 | 5445 ± 1402 | 1593 ± 441 |

### 2.2 Global chemical modification information and differential modification statistics

A total of 485 different chemical modifications were identified in 38 samples, of which 301 chemical modifications were identified in the children group, 329 chemical modifications were identified in the young group, and 106 chemical modifications were identified in the senior group. **Figure 1** shows the interaction diagram (Venn diagram) of modification in these three types of samples. For the specific content of the interaction, please refer to **Supplementary Table 1**.

**Fig 1.**
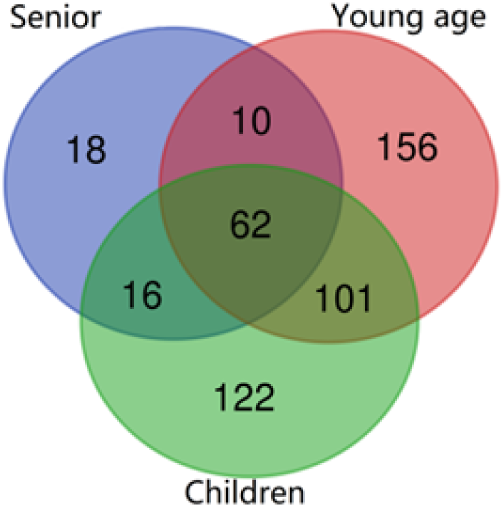
The Venn of different types of modification between three group samples

Global chemical modifications of 485 occurred at 20 amino acids and the N and C terminals. **Supplementary Table 2** shows the information on the modifications of each amino acid. As shown in the histogram of **Figure 2**, we calculated the ratio of amino acid modifications according to the ratio (modification sites number/ peptide number) of each modification in each age group. At the same time, we found that among the non-artificial modifications of proline(Pro, P), tryptophan(Trp, W), tyrosine(Tyr, Y), cysteine(Cys, C) and methionine(Met, M), oxidations and amino acid substitutions were dominant, especially in proline, tryptophan, and tyrosine, the oxidations rate of the senior is higher than that in other age groups, as shown in the pie chart in **Figure 2**. A total of 27 amino acids were substituted in the senior, of which there was no histidine substitution, and a total of 13 amino acids were replaced by histidine, which only occurred in children and young people, see **Supplementary Table 2** for details.

**Fig 2.**
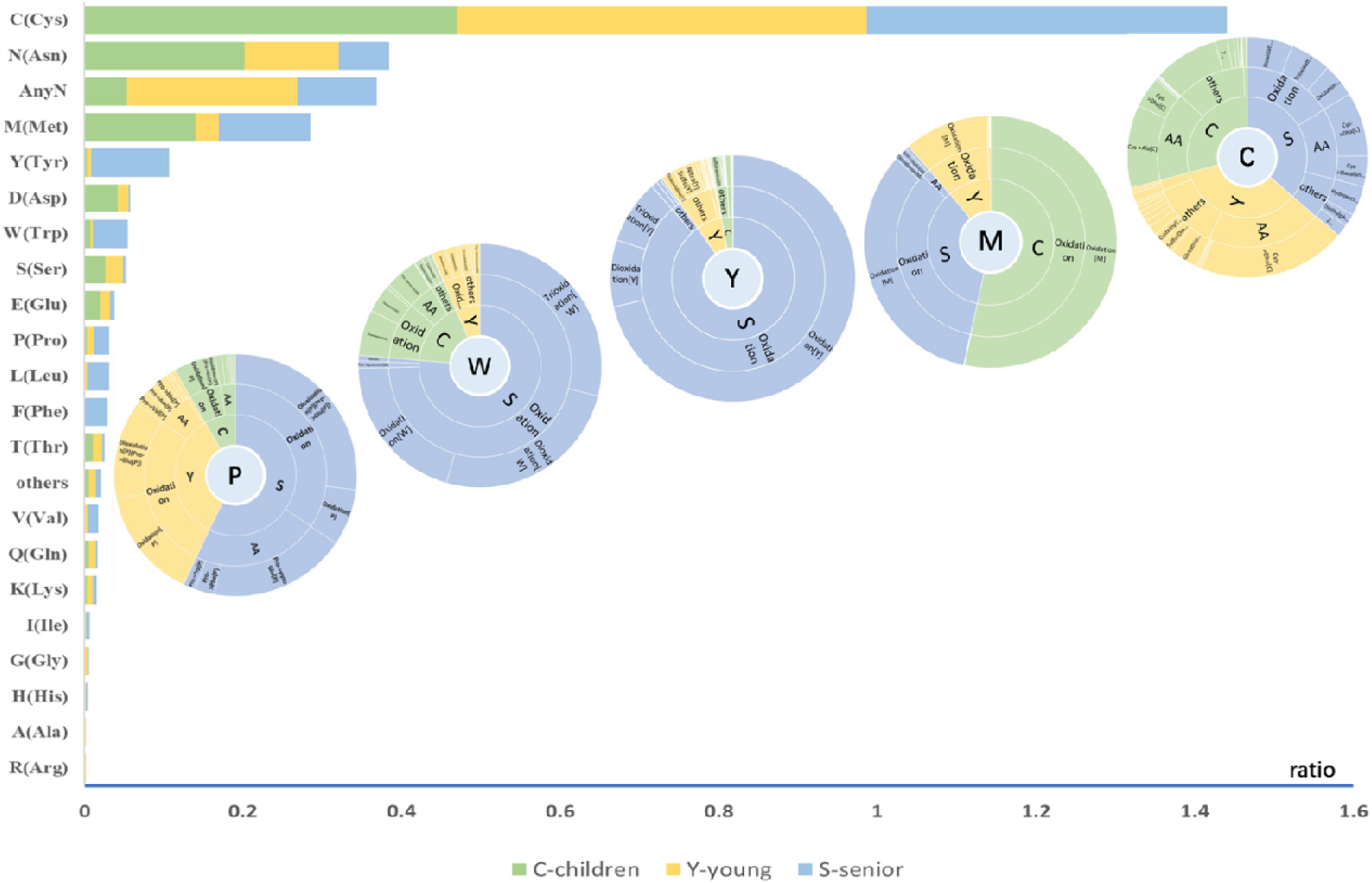
Statistics on the number and proportion of various amino acid site modifications between three age groups. The abscissa is the ratio of the number of various amino acid modification sites to the number of peptides identified. The pie chart showing the proportion of the same amino acid modification between different age groups, the detailed content of each part has been placed in the schedule

Unsupervised cluster analysis was performed on the global modifications and the 62 common modifications. It was found that the same sample of the senior group was abnormal and assigned to children and young groups in both heat maps. Besides, each sample was grouped into itself age group (**Figure S1** and **Figure S2**). There is a total of 40 modifications with statistical differences (p-value is less than 0.05), and the changes in each modification between different groups can be obtained through calculation and statistics. **Table 5** shows the details. Each differential modification was compared between different groups. The specific situation is shown in **Figure 3**. The significant differences between the 40 differential modifications were selected as follows: proline, tryptophan, tyrosine, cysteine, methionine oxidative modification, cysteine to dehydroalanine (Cys→Dha [C]), glycidamide adduct (glycidamide [AnyN-term]), phosphorylation of serine (Phospho [S]), the two protons of asparagine and glutamicacid replaced by calcium ions (Cation_Ca [E], Cation_Ca [D]), succinylation of the N-terminus of the protein (Succinyl [AnyN-term]), protein N-terminal carbamylation (Carbamyl [AnyN-term]), valine replaced by threonine (Val→Thr [V]), glutamicacid replaced by methionine (Glu→Met [E]), glutamine instead of glutamicacid (Glu→Gln [E]), convertion of glycosylated asparagine residues upon deglycosylation with N-glycosidase F (PNGase F) in H2O (Deamidated [N]), cysteine dehydrogenation (DiDehydro [C]).

**Fig 3.**
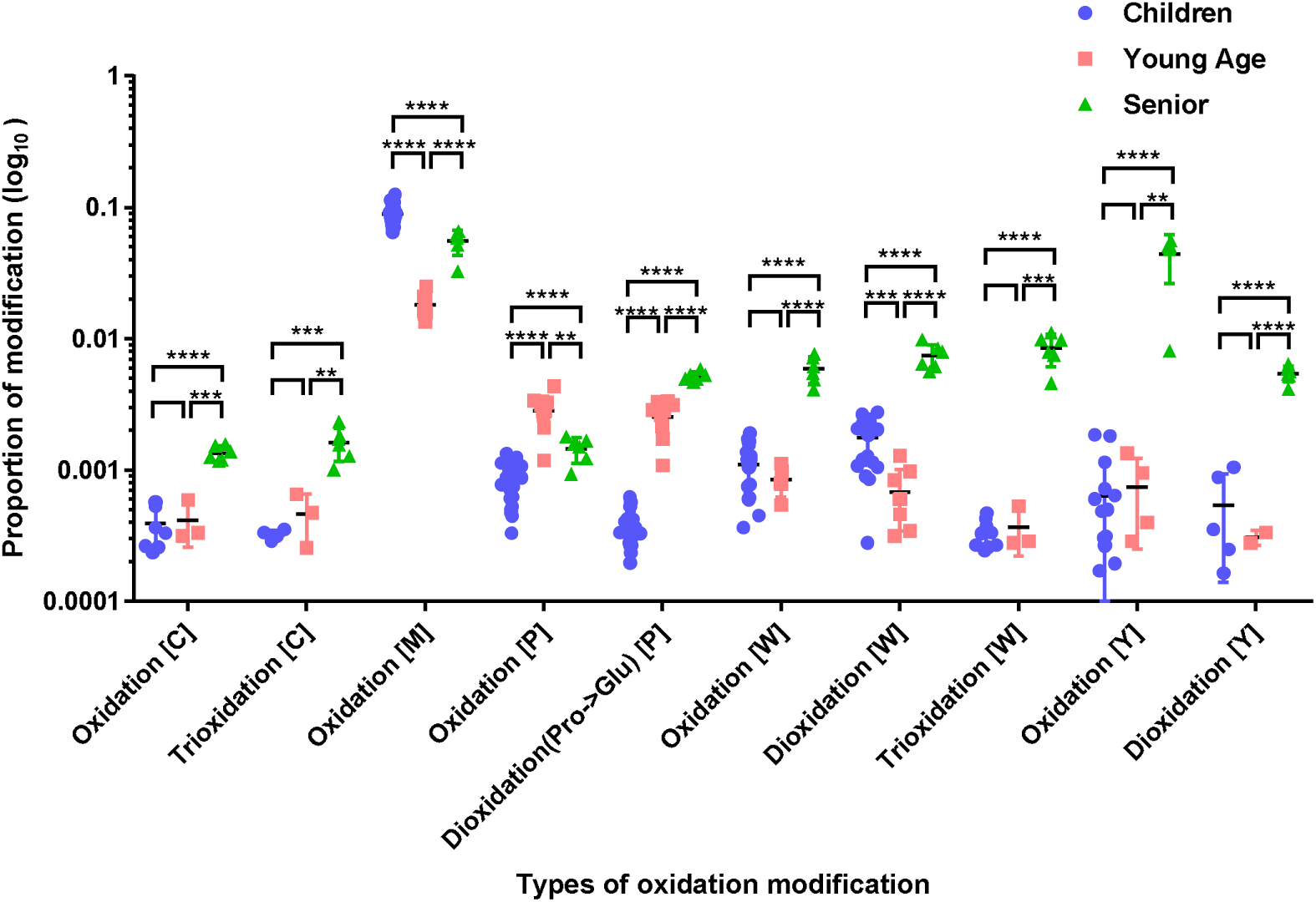
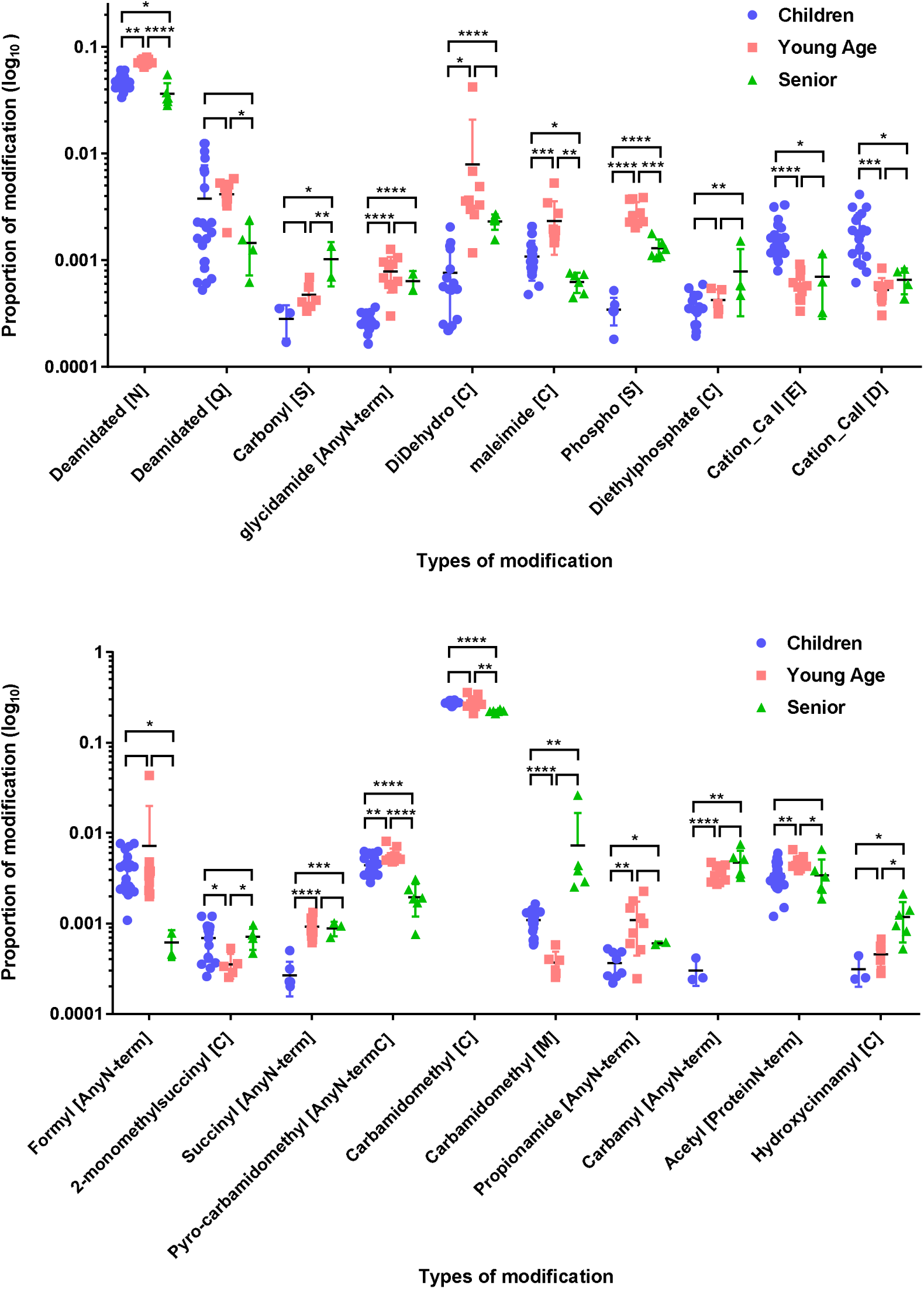
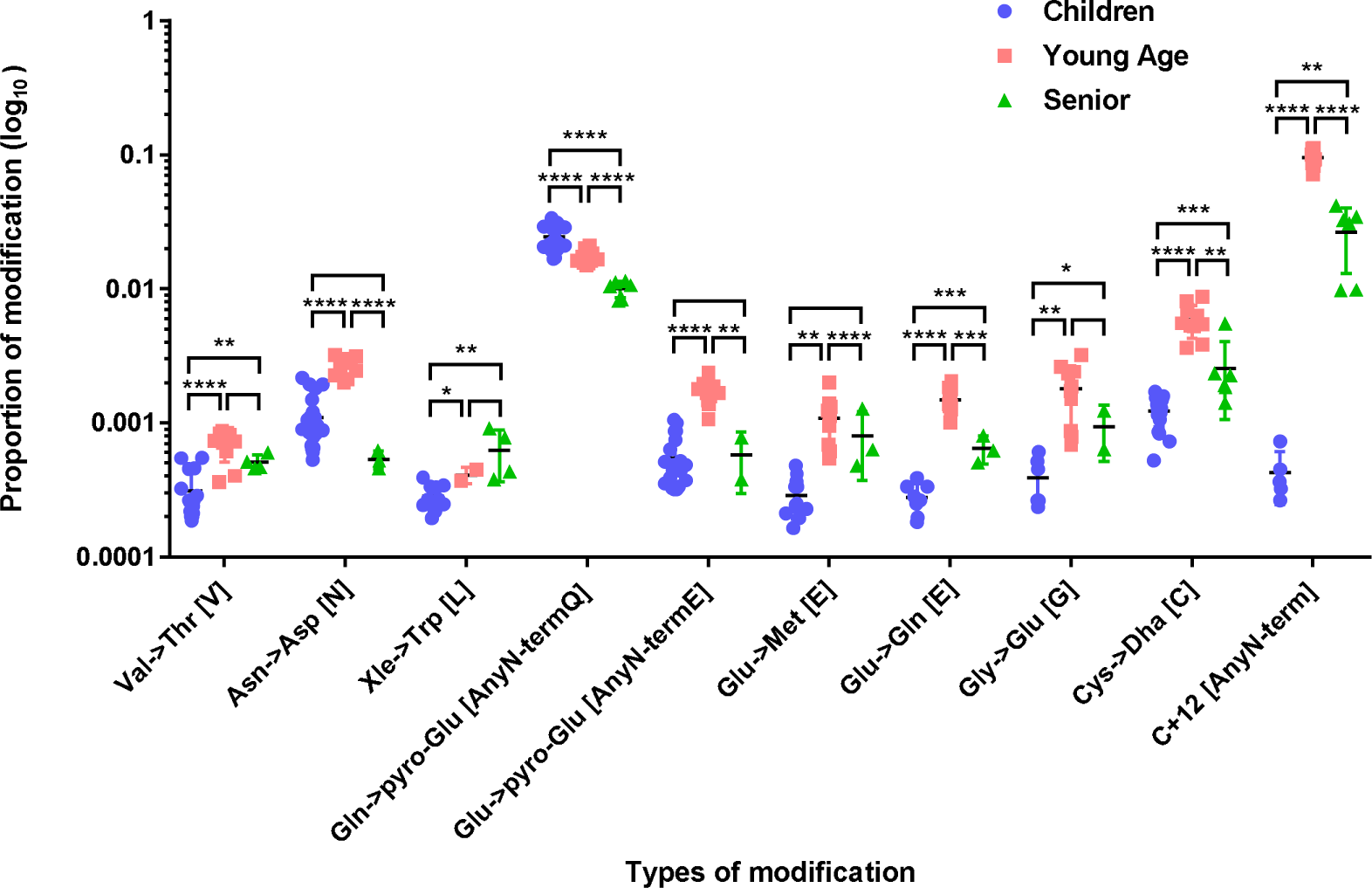
Comparison of modification differences among children group, young age, and senior group. The black horizontal line in the middle of the data represents the median, the upper and lower color lines represent the data quartile range, and the upper part of the data represents the significance between groups. As the number of labels increases, the significance increases. (* means p <0.05, ** means p <0.01, *** means p <0.005, **** means p <0.001)

**Table 5.** Differences and statistics of chemical modifications between different groups

| Modification Name | Classification | Senior/Children |  | Senior/Young |  | Children/Young |  |
| --- | --- | --- | --- | --- | --- | --- | --- |
|  |  | Fold change | P-value | Fold change | P-value | Fold change | P-value |
| Oxidation [C] | Post-translational | 3.46 ↑ | 9.73E-08 | 3.26 ↑ | 1.06E-04 | 1.06 ↓ | >0.05 |
| Trioxidation [C] | Chemical derivative | 5.04 ↑ | 5.52E-04 | 3.52 ↑ | 4.72E-03 | 1.43 ↓ | >0.05 |
| Oxidation [M] | Artefact | 1.61 ↓ | 3.38E-05 | 3.05 ↑ | 2.42E-07 | 4.93 ↑ | 6.29E-15 |
| Oxidation [P] | Post-translational | 1.79 ↑ | 5.88E-05 | 1.96 ↓ | 1.65E-03 | 3.57 ↓ | 1.97E-11 |
| Dioxidation(Pro→Glu) [P] | Post-translational | 14.11 ↑ | 2.99E-21 | 2.06 ↑ | 1.21E-06 | 6.67 ↓ | 2.32E-11 |
| Oxidation [W] | Artefact | 5.36 ↑ | 1.61E-13 | 6.99 ↑ | 1.93E-05 | 1.30 ↑ | >0.05 |
| Dioxidation [W] | Chemical derivative | 4.17 ↑ | 6.64E-13 | 10.94 ↑ | 7.10E-08 | 2.62 ↑ | 1.76E-04 |
| Trioxidation [W] | Chemical derivative | 1.76 ↑ | 8.35E-08 | 23.11 ↑ | 6.53E-04 | 1.11 ↓ | >0.05 |
| Oxidation [Y] | Post-translational | 69.11 ↑ | 6.96E-09 | 59.04 ↑ | 1.42E-03 | 1.18 ↓ | >0.05 |
| Dioxidation [Y] | Post-translational | 10.06 ↑ | 3.91E-07 | 17.67 ↑ | 9.79E-05 | 1.76 ↑ | >0.05 |
| Deamidated [N] | Artefact | 1.32 ↓ | 3.29E-03 | 2.00 ↓ | 7.29E-08 | 1.52 ↓ | 2.64E-11 |
| Deamidated [Q] | Artefact | 2.56 ↓ | >0.05 | 2.86 ↓ | 1.06E-08 | 1.10 ↓ | >0.05 |
| Carbonyl [S] | Chemical derivative | 3.64 ↑ | >0.05 | 2.15 ↑ | 1.41E-02 | 1.69 ↓ | 4.83E-02 |
| glycidamide [AnyN-term] | Chemical derivative | 2.41 ↑ | 1.84E-06 | 1.23 ↓ | >0.05 | 2.94 ↓ | 4.26E-07 |
| DiDehydro [C] | / | 3.03 ↑ | 4.16E-06 | 3.45 ↓ | >0.05 | 10.00 ↓ | 4.04E-02 |
| maleimide [C] | Chemical derivative | 1.72 ↓ | 2.01E-03 | 3.70 ↓ | 4.48E-03 | 2.17 ↓ | 2.04E-04 |
| Phospho [S] | Post-translational | 3.75 ↑ | 3.98E-06 | 2.17 ↓ | 1.74E-04 | 8.33 ↓ | 1.45E-07 |
| Diethylphosphate [C] | Chemical derivative | 2.19 ↑ | 4.61E-03 | 1.85 ↑ | >0.05 | 1.19 ↓ | >0.05 |
| Cation_Ca[II][E] | Artefact | 2.38 ↓ | 2.31E-02 | 1.19 ↑ | >0.05 | 2.80 ↑ | 2.24E-05 |
| Cation_Ca[II][D] | Artefact | 2.78 ↓ | 2.01E-02 | 1.24 ↑ | >0.05 | 3.48 ↑ | 2.68E-04 |
| Formyl [AnyN-term] | Post-translational | 6.67 ↓ | 2.17E-02 | 11.11 ↓ | >0.05 | 1.79 ↓ | >0.05 |
| 2-monomethylsuccinyl [C] | Chemical derivative | 1.04 ↑ | >0.05 | 2.02 ↑ | 1.07E-02 | 1.95 ↑ | 3.03E-02 |
| Succinyl [AnyN-term] | Post-translational | 3.30 ↑ | 1.01E-04 | 1.04 ↓ | >0.05 | 3.45 ↓ | 5.34E-06 |
| Pyro-carbamidomethyl [AnyN-termC] | Chemical derivative | 2.22 ↓ | 2.75E-05 | 2.86 ↓ | 4.27E-06 | 1.28 ↓ | 4.83E-03 |
| Carbamidomethyl [C] | Chemical derivative | 1.25 ↓ | 1.37E-09 | 1.27 ↓ | 6.71E-03 | 1.01 ↓ | >0.05 |
| Carbamidomethyl [M] | Chemical derivative | 6.68 ↑ | 2.86E-03 | 19.79 ↑ | >0.05 | 2.96 ↑ | 3.44E-06 |
| Propionamide [AnyN-term] | Chemical derivative | 1.65 ↑ | 3.08E-02 | 1.82 ↓ | >0.05 | 3.03 ↓ | 7.41E-03 |
| Carbamyl [AnyN-term] | Multiple | 15.65 ↑ | 2.55E-03 | 1.31 ↑ | >0.05 | 12.50 ↓ | 1.32E-05 |
| Acetyl [ProteinN-term] | Multiple | 1.03 ↓ | >0.05 | 1.39 ↓ | 4.85E-02 | 1.35 ↓ | 5.84E-03 |
| Hydroxycinnamyl [C] | Post-translational | 3.76 ↑ | 3.66E-02 | 2.58 ↑ | 1.21E-02 | 1.45 ↓ | >0.05 |
| Val→Thr [V] | AA substitution | 1.64 ↑ | 8.68E-03 | 1.35 ↓ | >0.05 | 2.22 ↓ | 2.55E-06 |
| Asn→Asp [N] | AA substitution | 2.04 ↓ | >0.05 | 5.00 ↓ | 4.64E-06 | 2.44 ↓ | 7.20E-10 |
| Xle→Trp [L] | AA substitution | 2.21 ↑ | 1.45E-03 | 1.54 ↑ | >0.05 | 1.43 ↓ | 2.71E-02 |
| Gln→pyro-Glu<br>[AnyN-termQ] | Artefact | 2.44 ↓ | 1.63E-07 | 1.72 ↓ | 2.64E-06 | 1.43 ↑ | 8.52E-05 |
| Glu→pyro-Glu<br>[AnyN-termE] | / | 1.06 ↑ | >0.05 | 2.94 ↓ | 2.01E-03 | 3.13 ↓ | 9.54E-12 |
| Glu→Met [E] | AA substitution | 2.77 ↑ | 1.08E-03 | 1.35 ↓ | >0.05 | 3.70 ↓ | 5.31E-06 |
| Glu→Gln [E] | AA substitution | 2.31 ↑ | 3.14E-04 | 2.33 ↓ | 9.41E-04 | 5.26 ↓ | 9.51E-09 |
| Gly→Glu [G] | AA substitution | 2.40 ↑ | 2.44E-02 | 1.92 ↓ | >0.05 | 4.55 ↓ | 1.41E-03 |
| Cys→Dha [C] | Chemical derivative | 2.07 ↑ | 3.90E-04 | 2.33 ↓ | 1.13E-03 | 4.76 ↓ | 6.30E-14 |
| C+12 [AnyN-term] | / | 62.20 ↑ | 2.02E-03 | 3.57 ↓ | 1.25E-07 | 100.0 ↓ | 1.15E-09 |

### 2.3 Overall oxidative modifications or amino acid substitutions are the main differences from children to senior people

Due to the significant difference in oxidative modification, and its close correlation with age, we found out all the oxidative modifications in 485 kinds of modifications, including oxidative modification of proline, tryptophan, tyrosine, cysteine, methionine and other insignificant and non-differential oxidative modifications (29 types in total). Unsupervised cluster analysis showed that it can be a perfect feature to distinguish these different age groups well, **Figure 4** shows the details. It is worth mentioning that all the amino acid substitutions (149 types in total) were found in the global modification at the same time. And cluster analysis showed that it could be an obvious feature to distinguish three different age groups very well, as shown in **Figure 5**.

**Fig 4.**
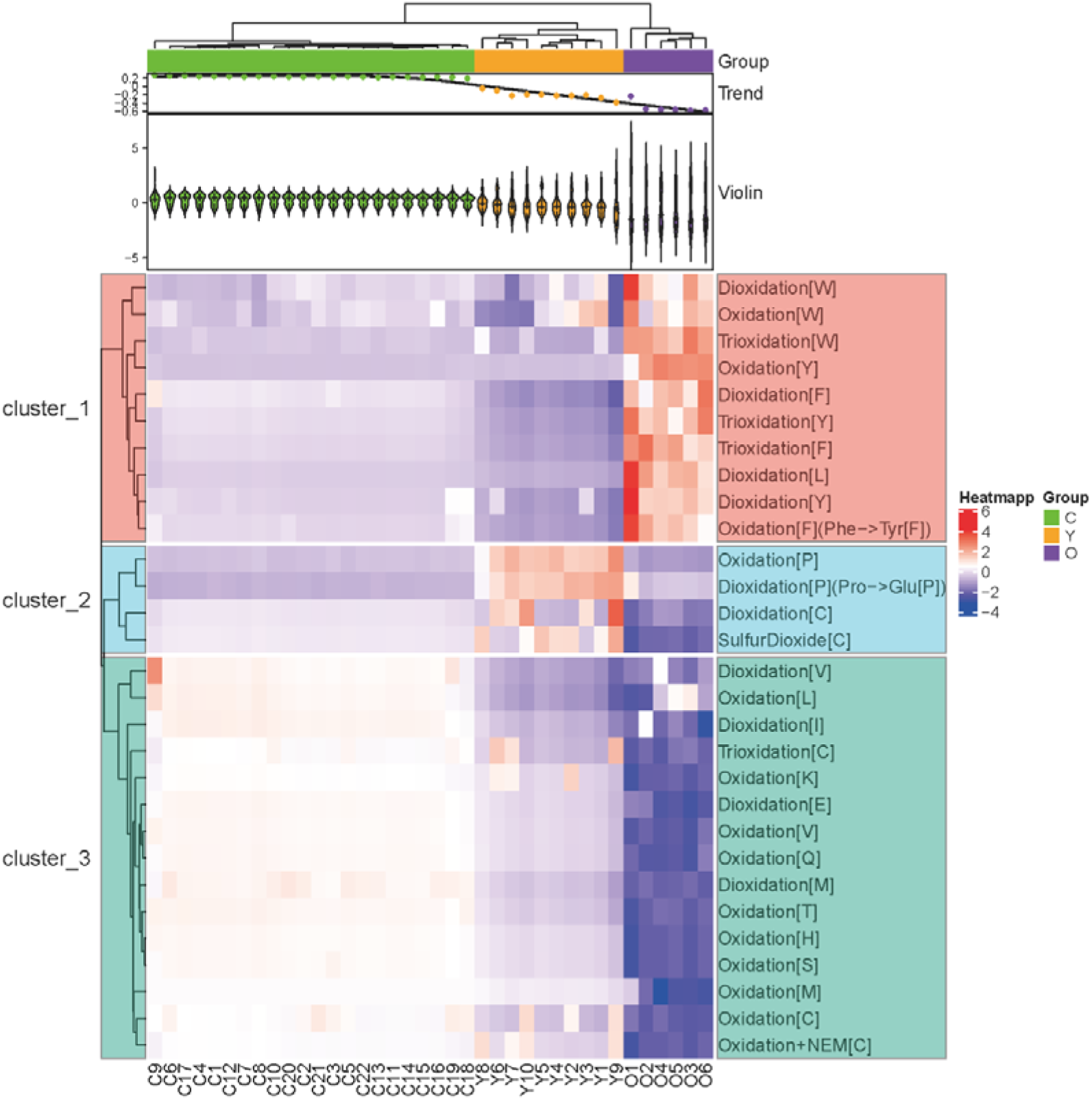
Cluster analysis results of all oxidative modifications (29 types) among the three groups of samples. Children group covers with green, and young age group covers with orange, the senior group covers with purple.

**Fig 5.**
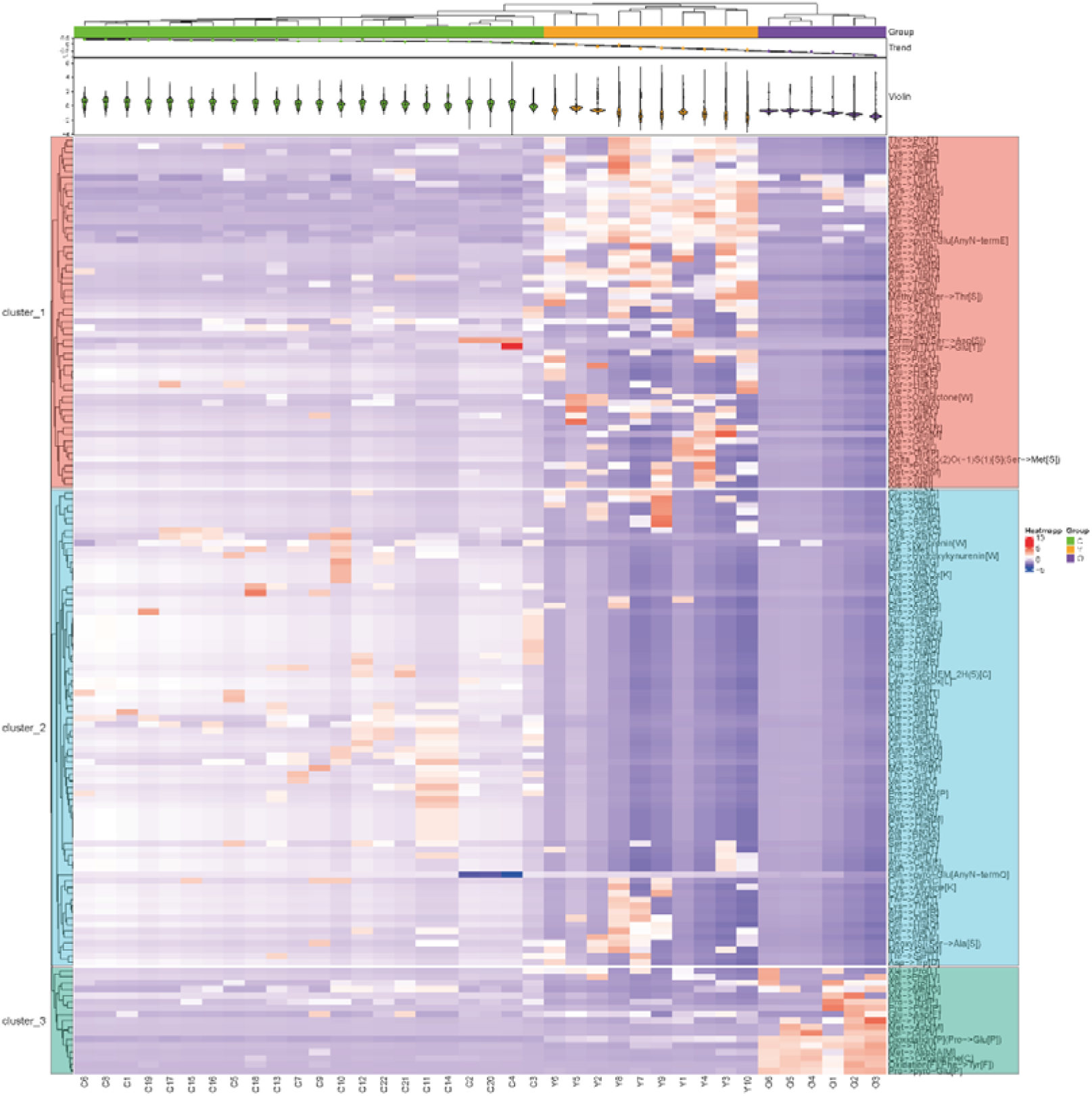
Cluster analysis results of all amino acid substitutions (149 types) among the three groups of samples. Children group covers with green, and young age group covers with orange, the senior group covers with purple.

Because the significantly differential oxidative modifications are concentrated in proline, tryptophan, tyrosine, and cysteine, we found that all these modifications (132 in total) for cluster analysis. It showed that except for one sample of the senior group, which was classified under the branch of the young group, the senior group could be well distinguished from the children and young groups (**Figure S3**). At the same time, all cysteine modifications (81 in total) and N-terminal modifications (61 in total) were analyzed by cluster analysis, as shown in **Figure S4** and **Figure S5**. Methylation and ethylation modifications are unique in children and young groups, while alkylation modifications are rarely found in the senior group.

### 2.4 Differential modifications in the senior group

The number of oxidative modification of various amino acids such as tryptophan (W), proline (P), cysteine (C), tyrosine (Y) and carbonylation of serine (S) in urine protein samples of the senior group is significantly more than that of children and young samples, and the carbamidomethyl (IAA-introduced alkylation modification) is significantly lower than that of children and young samples, because of reducing disulfide bonds, which indirectly shows that the disulfide bonds integrity in the senior group is significantly lower than that of young group and children group. The phosphorylation of serine, maleylation of cysteine, aspartic acid (D) replaced by asparagine (N), methionine (M) and glutamine (Q) replaced by glutamic acid (E), glycine (G) replaced by glutamic acid(E) and cysteine dethiolation to dehydroalanine (Dha) were significantly higher in the young group than those in children and senior group. In the children’s group, calcium ions-related modifications of glutamic acid and aspartic acid are significantly higher than that of the young group and the senior group, but the number of modifications involving the dehydrogenation of cysteine and the modification of N-terminus, such as carbamoylation, glycamylation, succinyl is lower than that of the young group and the senior group. Also, the modification of glutamine to pyro-Glu shows a decreasing trend with aging.

Oxidation of isoleucine (I), glutamine(Q), glutamic acid(E), phenylalanine (F), and phosphorylation of aspartic acid, glutamic acid, and tyrosine are unique modifications in the senior group (see **Supplementary Table 3** for more information). **Figure 6** shows the modification identification information of several peptides in the secondary mass spectrometry (MS2) results of the senior group.

**Fig 6.**
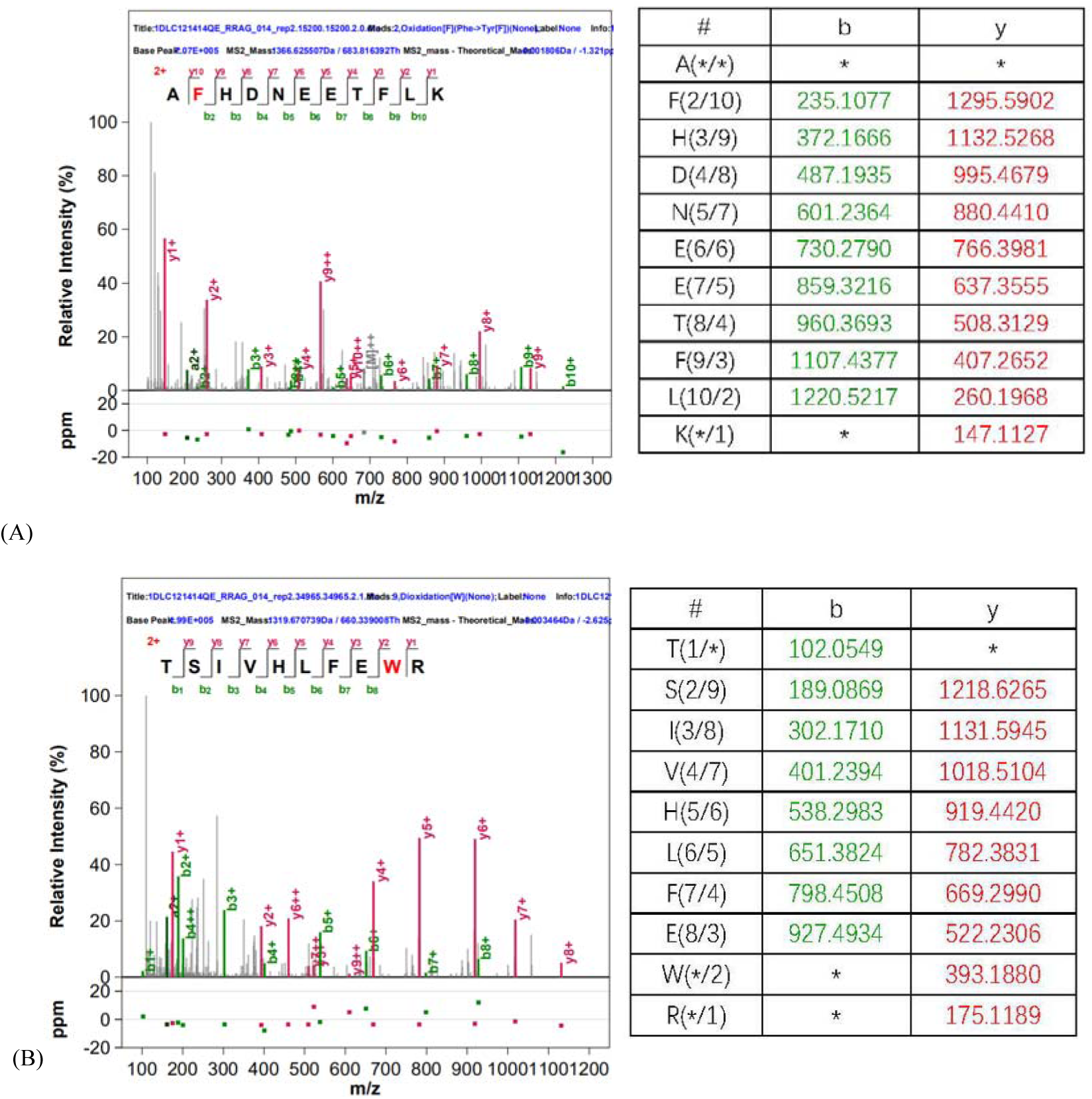
Modified identification results of several peptides by MS in samples from the senior group. (A) Alpha-amylase peptide in the tryptophan (Trp) site dioxidation b / y ion information; (B) serum albumin peptide the oxidation b / y ion information of phenylalanine (Phe) oxidized to tyrosine (Tyr);

### 2.5 Oxidation ratio of the senior group is the highest and PPI analysis

Randomly selecting one sample in each group to find the proteins where the above modifications are located, and take a union obtaining 971 proteins, and perform protein interaction analysis (PPI BIOGRID—Network Topology-based Analysis, NTA) at WebGestalt (http://www.webgestalt.org), running under default parameters. The analysis of related biological process pathways shows that these modifications affect the function of the body’s immune system and coagulation-related system (**Figure 7**). The oxidative modifications identified above exist in 446 proteins, accounting for 45.9% of the total. Oxidation of methionine may be interfered during the experiment. Some modifications in the UniMod database were determined to be artificially introduced, and non-artificial modifications exist in 359 proteins. And oxidative modification of proline, tryptophan, tyrosine, and cysteine existed in 256 proteins, accounting for 71.3%. By looking up the statistics of the non-artificial modified peptide in Peptide.all_result of pBuild, and directly finding the number of non-artificial modification sites in pFind.Summary, through comparison and calculation, it was found that oxidative modifications ratio of children group samples was 12.97% (in terms of peptides, the same below), the oxidation modification ratio of the young group was 13.71%, and the oxidation modification ratio of the senior group was as high as 29.76%, which was significantly higher than that of children group and the young group. For details, see **Table 6**.

**Fig 7.**
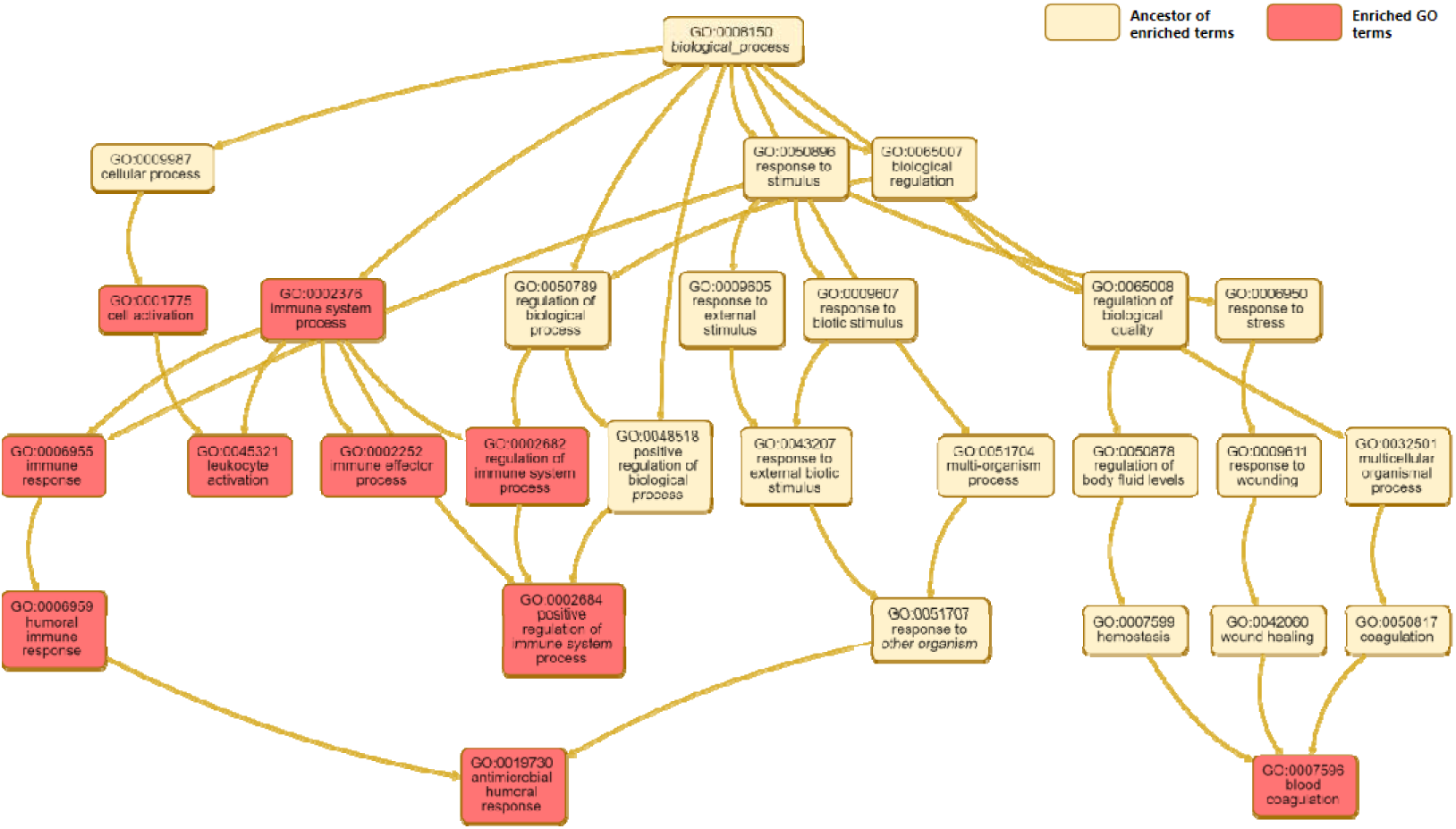
Enriched GO terms graph of modified proteins in three age groups.

**Table 6.**
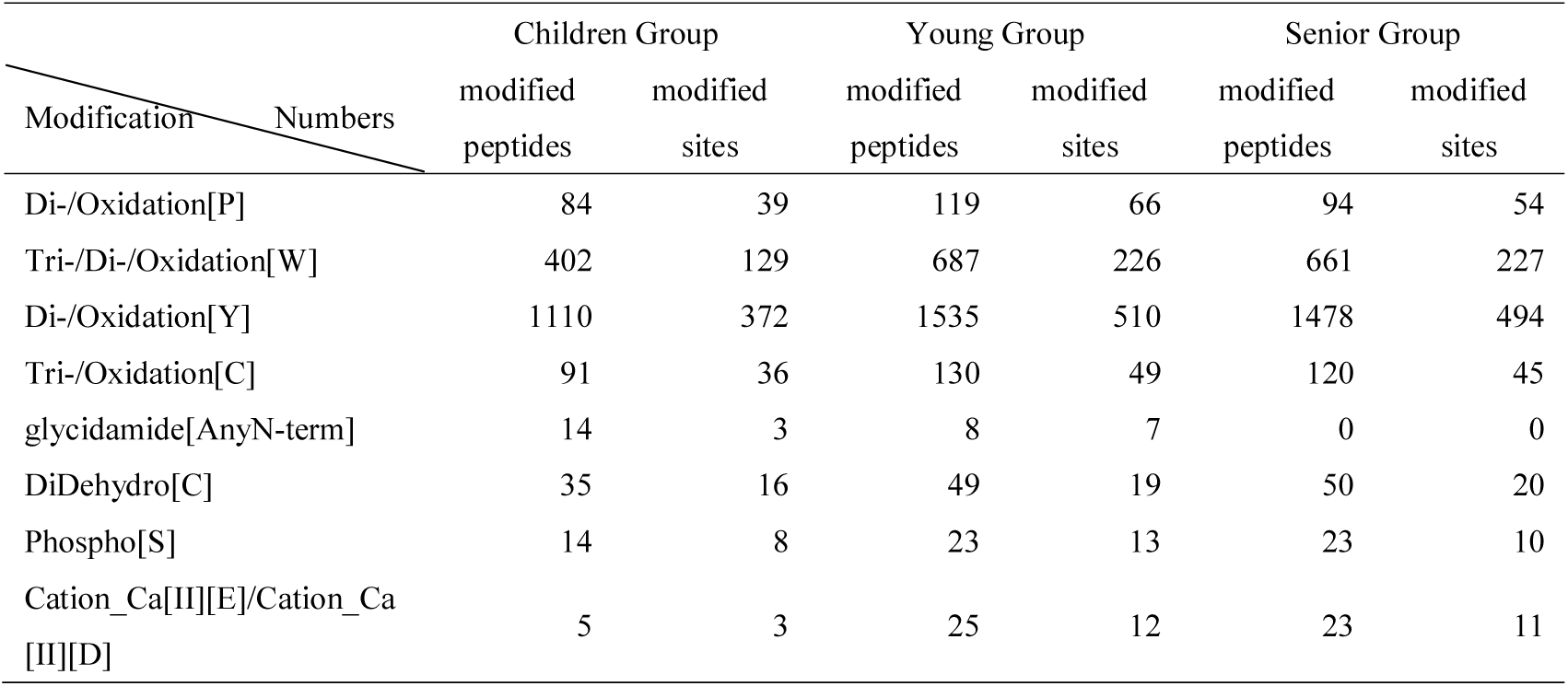
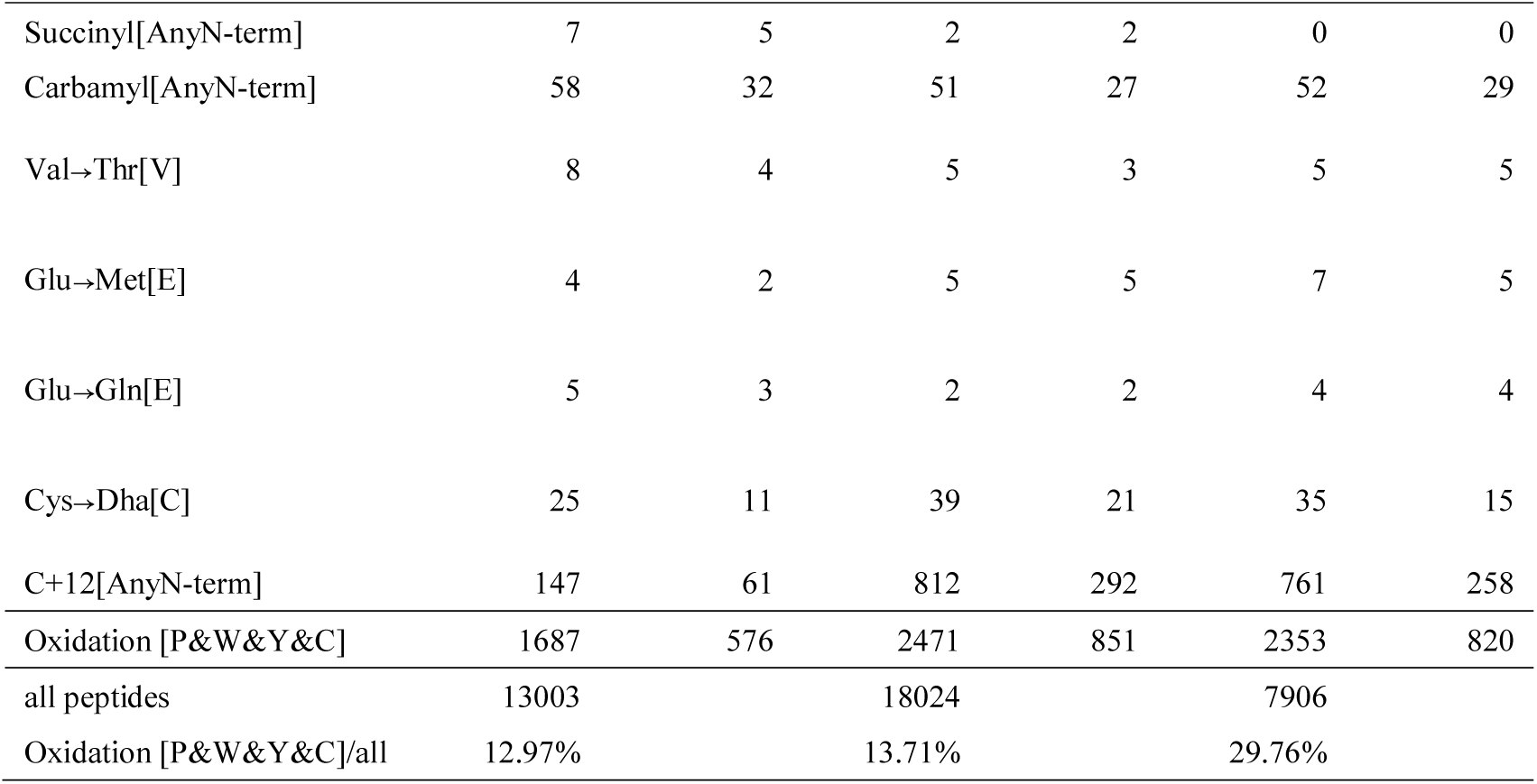
Statistics of non-artificial modified peptides and sites in three age groups samples

### 2.6 Random grouping test of 10 oxidative modifications

Ten oxidative modifications that meet the screening criteria of difference (p-value less than 0.01, fold change greater than 2 or less than 0.5) were randomly and independently grouped to verify the false positive rate of each oxidative modification. All the data of 38 all samples (children and young people as a group, 32 samples, the elderly as a group, 6 samples) were randomly divided into two groups with a total of 2,760,681 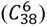 different combinations. Through the same differential screening conditions, the combination under each condition was statistically analyzed. After detailed calculation and statistics, random combinations of ten oxidative modifications were obtained (**Table 7**). Through the random grouping test, it is found that the randomness of 10 oxidative modifications is in the range of 0.005∼0.02, and the credibility is in the range of 97.9%∼99.5%. It has been proved that the difference of 10 oxidative modifications in different age groups is unlikely to be generated randomly.

**Table 7.** Random grouping calculation results in 10 oxidative modifications in urine proteome

| Modification Name | Total number of occurrences | Occurrence frequency | Credibility |
| --- | --- | --- | --- |
| Oxidation[M] | 16828 | 0.006096 | 99.39 % |
| Oxidation[P] | 14163 | 0.00513 | 99.49 % |
| Dioxidation[W] | 37362 | 0.013534 | 98.65 % |
| Oxidation[W] | 39456 | 0.014292 | 98.57 % |
| Dioxidation[P](Pro->Glu[P]) | 25512 | 0.009241 | 99.08 % |
| Oxidation[Y] | 57233 | 0.020731 | 97.93 % |
| Trioxidation[W] | 46243 | 0.016751 | 98.32 % |
| Oxidation[C] | 32649 | 0.011826 | 98.82 % |
| Dioxidation[Y] | 46421 | 0.016815 | 98.32 % |
| Trioxidation[C] | 42133 | 0.015262 | 98.47 % |

## 3. Discussion

Our study found that the oxidation degree of various protein sites of in the senior group was significantly higher than that of the children and the young group. It can be inferred that urine is the main way to eliminate the production of oxidative and aging proteins in the human body. The amount of oxidized proteins produced by the senior was very large so that the amount of oxidized proteins excluded was also very large. The oxidized ability of a protein is directly related to the characteristics of its primary structure and three-dimensional structure, which may lead to the destruction of its structure and function [21, 22]. An increase in the level of oxidatively modified proteins has been reported in senior erythrocytes of higher density and cultured human fibroblasts from normal senior donors and individuals suffering from progeria and Werner’s syndrome [23, 24].The degree to which oxidized proteins accumulated in the human body is an important manifestation of human aging, which can be used to judge the degree of human aging and whether anti-aging drugs are effective.

Among the three age groups proline oxidative modified peptides, we found that they belong to Collagen (A0A384MDU2_HUMAN) and Collagen alpha-1(I) chain preproprotein (H9C5C5_HUMAN). Proline and hydroxyproline account for approximately 1/4-1/5 of collagen amino acids. Collagen is the main component of the degraded extracellular matrix, which provides amino acids and energy for cell growth and proliferation, and it is also found that collagen degrades in the process of inflammation and tumour progression development [25, 26]. The overexpression of proline dehydrogenase in some cancer cell lines is related to apoptosis, and the mechanism includes the oxidation of proline to produce Reactive Oxygen Species (ROS) in cells [27, 28]. Because the overexpression of proline dehydrogenase is also associated with the reduction of tumor formation in vivo, it is considered to be a direct regulator of p53-induced tumor suppression [28, 29]. Moreover, some cancer cells may utilize proline oxidation to promote proliferation and resist stress [30–32]. It is inferred that the rapid aging of collagen in the senior brings a large amount of oxidized proline-containing protein. These oxidized proteins are continuously discharged into the urine, so the content of oxidative proline in urine is obviously higher than that of young age and children.

We found that among the three age groups, some peptides were modified by tryptophan, tyrosine oxidation, and methionine instead of glutamic acid belong to the Fibrinogen alpha chain (FIBA_HUMAN), and we also found fiber proprotein peptides in the unique modification of the senior. Fibrinogen, one of the most abundant blood plasma proteins, is well known to be the most frequent target of PTM. It is the last member of the blood coagulation cascade reaction [33]. After being activated by thrombin [34–36], fibrinogen is transformed into fibrin and then polymerized into a complex fibrin network [37].The fibrin net, along with platelets, red blood cells, and some white blood cells is the main components of thrombosis [38, 39]. Under physiological conditions, thrombus can prevent blood loss at the injured sites, but under pathophysiological conditions, it can adhere to blood vessels (thrombosis) and be released into the blood (embolism), which may lead to stroke, myocardial infarction and so on [40, 41]. Oxidation produced during inflammation (e.g., oxidation of Met476 located in the αC domain) has been proven to cause fibrin fibers to become thinner, resulting in denser blood clots, more difficult to be proteolyzed, and deep vein thrombosis and increase pulmonary embolism risks [36].

Most of the evidence obtained from the data indicates that the oxidative modification of fibrinogen caused by increased oxidative stress may be related to disease pathogenesis [42]. Plasma fibrin clots consist of dense reticulated fibers. Because the diffusion of plasminogen in the clot is blocked, it is difficult to cleave [43], which can be observed in patients with cardiovascular diseases, chronic inflammation [44], liver diseases [45, 46], diabetes complications [47].

Tryptophan oxidation products accumulate with age and inflammation, while its decomposition products such as kynurenine causing osteoporosis and its oxidative metabolites may have different effects on the tissues where they accumulate [48]. The release of dual tyrosine, one of the oxidation products of tyrosine, only occurs under oxidative stress (exposure to H_2_O_2_) and subsequent proteolysis, so it can be considered as a special sign (unique indicators) of some diseases, such as atherosclerosis, acute inflammation, systemic bacterial infections and cataracts [1,49–51]. The oxidation of histidine, tyrosine, and tryptophan is a reaction to various reactive oxygen species. Although these modifications are generally considered irreversible, the correlation between them and the regulation of biological proteins is still unclear [52, 53].

The most common reversible redox modification sites in proteins include redox-active transition metal ion centres (e.g., heme groups, iron-sulfur protein centres, and zinc-sulfur protein centres) and oxidation sensitive amino acid side chains (mainly cysteine, selenocysteine and methionine residues). Their oxidative modification can enhance or inhibit enzyme activity (e.g., by oxidizing cysteine with catalytic activity), but can also lead to changes in protein-protein interactions, subcellular transport, protein transformation, etc [52]. The research field of redox signals mainly focuses on the reversible oxidation of highly conserved cysteine residues in proteins, and its relative abundance is increasing with the evolution of multicellular organisms [54, 55]. Compared with the rapidly growing literature on cysteine oxidation, methionine oxidation as a potential signal mechanism has not been fully understood. To a great extent, this gap in knowledge is due to the lack of biochemical reagents for evaluating methionine oxidation in biological systems, and only the mass spectrometry (MS) method can successfully prove methionine sulfoxidation. The sulfoxidation of methionine residues significantly improves the hydrophilicity of methionine, which can significantly change the physical and chemical properties of proteins, thus enhancing or inhibiting the activity of proteins [56]. Similar to reversible cysteine oxidation, it has been found that this enzyme system can enhance and reverse the methionine sulfoxidation [57].

Recent proteomic analysis showed that methionine residues around phosphorylation sites were preferentially oxidized in vivo under stress conditions, and there was a complex relationship between methionine oxidation and protein phosphorylation pathway [56]. Aging is associated with the gradual oxidation of the extracellular environment. The redox state of human plasma is defined by the concentrations of cysteine and cystine, and the oxidation degree becomes higher and higher with the increase of age [58]. Direct oxidation of cysteine and methionine (uncommon) residues is a major reaction; this is usually faster than H_2_O_2_-mediated oxidation, resulting in changes in protein activity and function. Unlike H_2_O_2_ can be rapidly removed by protective enzymes, while protein peroxides can only be slowly removed, and catabolism is the main destination. Although proteases and lysosomal enzymes and other proteases (such as mitochondrial Lon) are effective for the turnover of modified proteins, protein hydroperoxides inhibit these pathways, which may contribute to the accumulation of modified proteins in cells. Existing evidence supports the connection between protein oxidation and various human pathologies, but it remains to be determined whether the connection is causal [59]. Methionine oxidation may be an important endogenous mechanism to regulate transient receptor potential vanilloid 2 (TRPV2) activity, and its co-expression plays a key role in phagocytosis of giant cells [60].

Iodoacetamide alkylation of cysteine site (carbamidomethyl) reflects the amount of sulfhydryl groups in the protein chains. The sulfhydryl group in cysteine is an important factor involved in the formation of protein disulfide bonds, and the amount of disulfide bonds can reflect the integrity of the protein spatial structure. Therefore, the integrity of the protein structure can be known by comparing the carbamidomethyl in urine samples of different ages. We found that there was no significant difference between the young group and the children group in cysteine carbamidomethyl. The cysteine carbamidomethyl in the senior group was significantly lower than that of the young group and the children group, which indicated that the damage degree of protein spatial structures increased with the increase of age.

Except for oxidative modification, we also explored the replacement of cysteine by dehydroalanine and the carbamylation of the protein N-terminus in the previous article “Why are there proteins in the urine of healthy people?”[61]. Studies have shown that in blood samples of patients with chronic liver disease, chronic kidney disease, or diabetes, cysteine Cys34 modification in human serum albumin (HSA) can be used as a marker for oxidative stress-related diseases [62]. The conversion of cysteine to dehydroalanine (Dha) is irreversible. Dha is an electrophilic reagent, which has an unsaturated carbon-carbon double bond in the Dha-containing peptide. It can be used as an alkylating agent for the protein chain in vitro [63], and many peptides containing dehydroalanine have certain toxicity [64]. Therefore, the modified proteins or peptide are also toxic to some extent. We previously speculated that if they stay in the body for a long time, they will have toxic effects on the body, and it is better to immediately excrete them. Thus, a large number of proteins can be found in urine to be modified into dehydroalanine will. Carbamylation is also an irreversible non-enzymatic modification process, and its formation process is that the decomposition products of urea react with the N-terminus of the protein or the side chain of lysine residues, which is related to protein aging [65]. Lysine carbamylation can promote the coordination of metal ions to specific enzyme activities [66, 67]. It has been reported that the plasma carbamylation levels of patients with elevated urea levels (such as nephropathy) are significantly increased [68].

Glycidamide (GA) is a DNA-reactive metabolite of acrylamide (AA), and AA is recognized as one of the carcinogenic dietary factors. Its carcinogenicity is related to the ability of glycine forming a DNA adduct, that is, modification at the N-terminus of the protein [69–71].

The complex developmental procedures and the regulation of signalling pathways among many cells in vertebrate species strongly depend on phosphorylation-mediated signalling pathways [72–77]. The phosphorylation of serine and threonine is more likely to be non-functional in disordered areas. In a study of 2018, based on large-scale data of human and mouse phosphorylation sites, it was found that the sites in the senior group were more likely to function and participate in the signaling pathways than those in the young group, and serine phosphorylation may be related to the occurrence of neurodegenerative diseases [78]. Mutations of some specific genes lead to neurodegeneration through the loss of protein function [79–81], PTM can regulate the structure and function of proteins, and also regulate toxicity among several pathogenic proteins. Recent studies have confirmed the relationship between phosphorylation and arginine methylation, and there is evidence that these modifications play an important role in neurodegeneration. For example, the decrease of Huntingtin (Htt) phosphorylation at serine 421 indicates that protein kinase B (Akt) signal dysregulation is involved in the pathogenesis [82, 83].

In our previous study, we found that in the plasma samples, the cysteine succinylation in the senior group was significantly different from that in the young group. In this study, it was also found that the N-terminus succinylation of the protein was significantly different. Although the study of succinylation is still in its infancy, and succinylation is the key to cell integration, the data clearly shows that succinylation has extensive effects on health and disease, and provides a coupling between metabolism and protein function in the nervous system and neurological diseases [84]. Carbamylation is an irreversible process in which the decomposition products of urea and isocyanic acid react with the N-terminus of the protein or the side chains of lysine and arginine residues, and of the proteins non-enzymatically modified [85–88].

The substitution of valine by threonine and glutamic acid by methionine occurred in the misacylation of the tRNA and editing of the charged tRNA [89, 90]. Previous studies have shown that natural glutamine (Gln) replacing glutamic acid (Glu) at the 3rd amino acid position of oxyntomodulin (OXM) can reduce the activity of glucagon receptor (GCGr) without affecting the activity of glucagon-like peptide 1 receptor (GLP-1r) [91].

Among the non-artificial modifications with significant differences, the oxidative modification account for a large portion. Bioinformatics enrichment analysis shows that most proteins are related to the immune process, leukocyte activation and blood coagulation. In previous studies, it was found that ribosomal proteins and proteins related to energy metabolism, including proteins related to the tricarboxylic acid cycle (TCA), mitochondrial respiration, and glycolysis, were inefficiently expressed in the muscles of the senior. Proteins related to innate immunity, adaptive immunity, protein stability, and alternative splicing were overexpressed [92, 93]. Aging is a complicated phenomenon. Aging itself is the foundation of the development of age-related diseases, such as cancer, neurodegenerative diseases, and type 2 diabetes. In recent years, different theories have been put forward in scientific research to explain the aging process. Up to now, there is no single theory that can completely explain all aspects of aging. Judging from the large amount of evidence discovered over the years, the damage accumulation theory is one of the most widely accepted where the accumulation of damage is thought to be caused by oxidative stress, and oxidative stress promotes protein modification and senescent cells how to respond to them through protein stabilization mechanisms, including antioxidant enzymes and proteolytic systems [94–97].

Protein oxidation is associated with many diseases, especially those related to aging. There is abundant evidence that oxidation exists in neurodegenerative diseases, such as Alzheimer’s disease (AD) [95,98–106], and Parkinson’s disease [107], protein carbonylation has been confirmed in the whole brain of AD patients [108, 109], Lewy body dementia patients [110] and Parkinson’s disease patients [111]. Besides, protein carbonyls are also present in various diseases such as acute/adult respiratory distress syndrome [112, 113], chronic lung disease [114–117], amyotrophic lateral sclerosis [118, 119], rheumatoid arthritis and juvenile chronic arthritis [120, 121], severe sepsis [122, 123], cystic fibrosis [124, 125], cataractogenesis [126], age-related macular degeneration [127], chronic renal failure, uremia [128–131], Diabetes mellitus Type I and Type II [132–136], inflammatory bowel disease [134], ischemia-reperfusion [137], systemic amyloidosis [138], and essential arterial hypertension [139]. Protein oxidation is more obvious in age-related diseases. The above shows many protein oxidation diseases that have been confirmed so far. Protein oxidation may be the cause or consequence of these diseases, which affect almost all organ systems.

Aging is a predominant risk factor for several chronic diseases that limit healthspan [140]. Therefore, the aging mechanism is increasingly considered as potential therapeutic targets [16]. PTMs exist naturally in the human body and participate in many physiological functions such as cell differentiation and gene regulation. However, at high concentrations, they may indicate serious diseases, such as myocardial infarction, venous thromboembolism, arterial and venous thrombosis, pulmonary embolism, and cancer [141–145], and emerge as important markers of aging and aging-related diseases [146, 147]. Compared with blood, urine is non-invasive and easy to collect. What counts is urine formation is a process of accumulation, in which changes in the abundance and post-translational modification of proteins and accumulation of some covalently modified proteins maybe emerge earlier to represent hallmarks of biological aging [148].

In recent years, with the aggravation of elderly populations, the incidence of age-related chronic diseases is becoming prevalent, which will bring unprecedented challenges to individuals, societies, and their health care systems. Therefore, there is an urgent need to understand physiological aging and achieve “healthy” aging by reducing age-related pathological changes [149]. Recent studies have found that damaged dysfunctional or toxic proteins and silent mutations are the phenotypes of aging, and may also be the causes of related diseases [150]. Among them, the irreversible oxidative modifications of proteins are becoming more and more prominent in key cellular pathways and the pathogenesis of age-related diseases [151, 152].

We compared the global modifications of three different age groups. The overall oxidative modifications and all amino acid substitutions are obvious features to distinguish different age groups perfectly. Our previous research speculated that many amino acids lost their original function due to modification or substitution, and then excreted into the urine. These changes may be related to aging and even some pathological changes in the body. Therefore, the global modifications of the urinary proteome may have some fresh and significant implications for exploring the aging mechanism and searching for early biomarkers of age-related diseases.

## Acknowledgment

The authors thank the research of Vitko D, Cho PS, Kostel SA, et al. and Yu Y, Sikorski P, Smith M, et al. Their available data helps us to draw the conclusion. We thank Dr. Hao Chi (Institute of Computing Technology, CAS) for help in using pFind, and thank Dr. Shisheng Wang (West China Hospital, Sichuan University) and Dr. Chengpin Shen (Omicsolution Co., Ltd) for giving some advice about data analysis and ‘Wu Kong’ platform (https://www.omicsolution.org/wkomics/main/) for relative cluster analysis.

---

**Fig S1.**
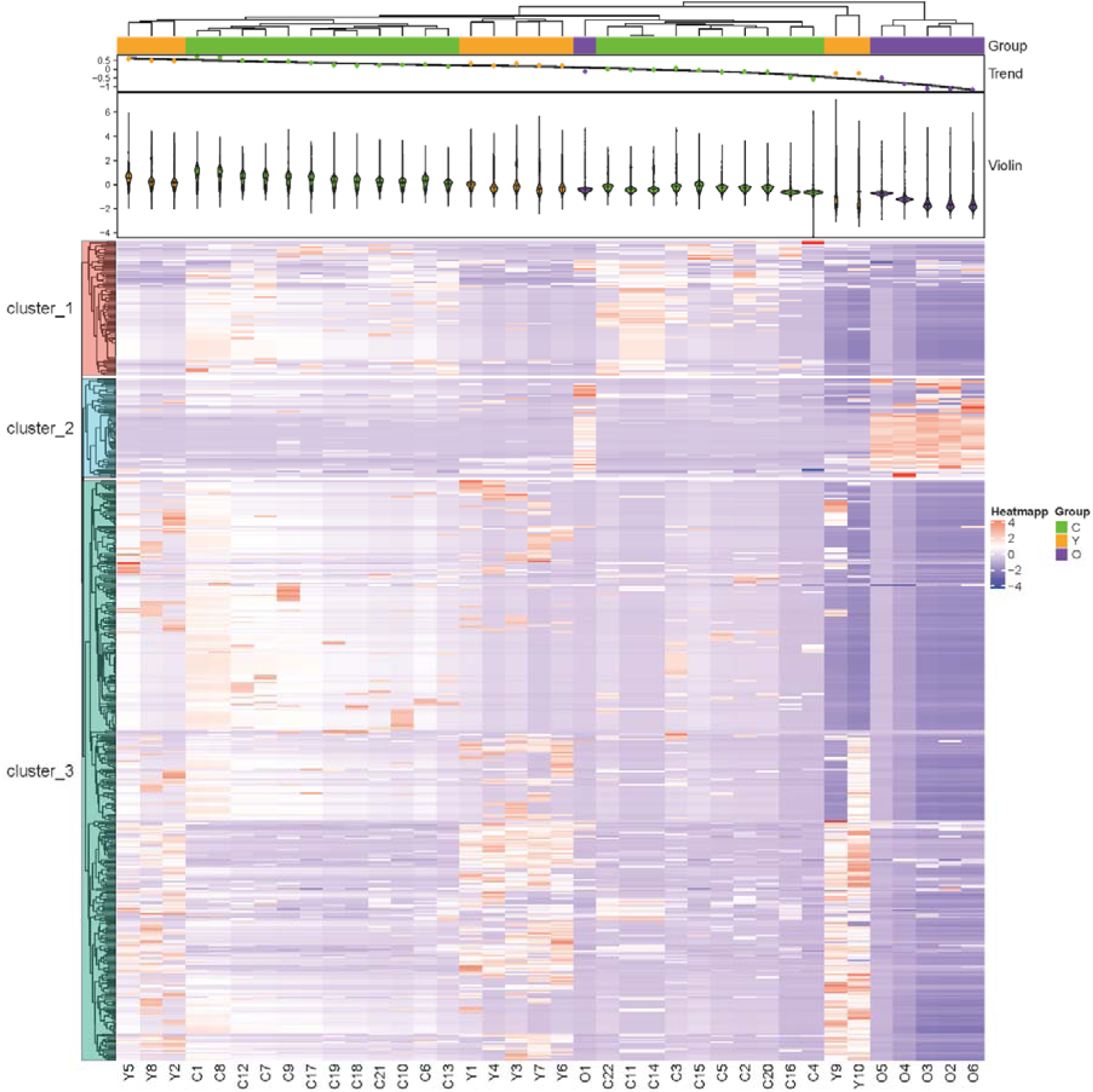
Cluster analysis results of the total modifications for three groups samples. Children group covers with green, and young age group covers with orange, the senior group covers with purple.

**Fig S2.**
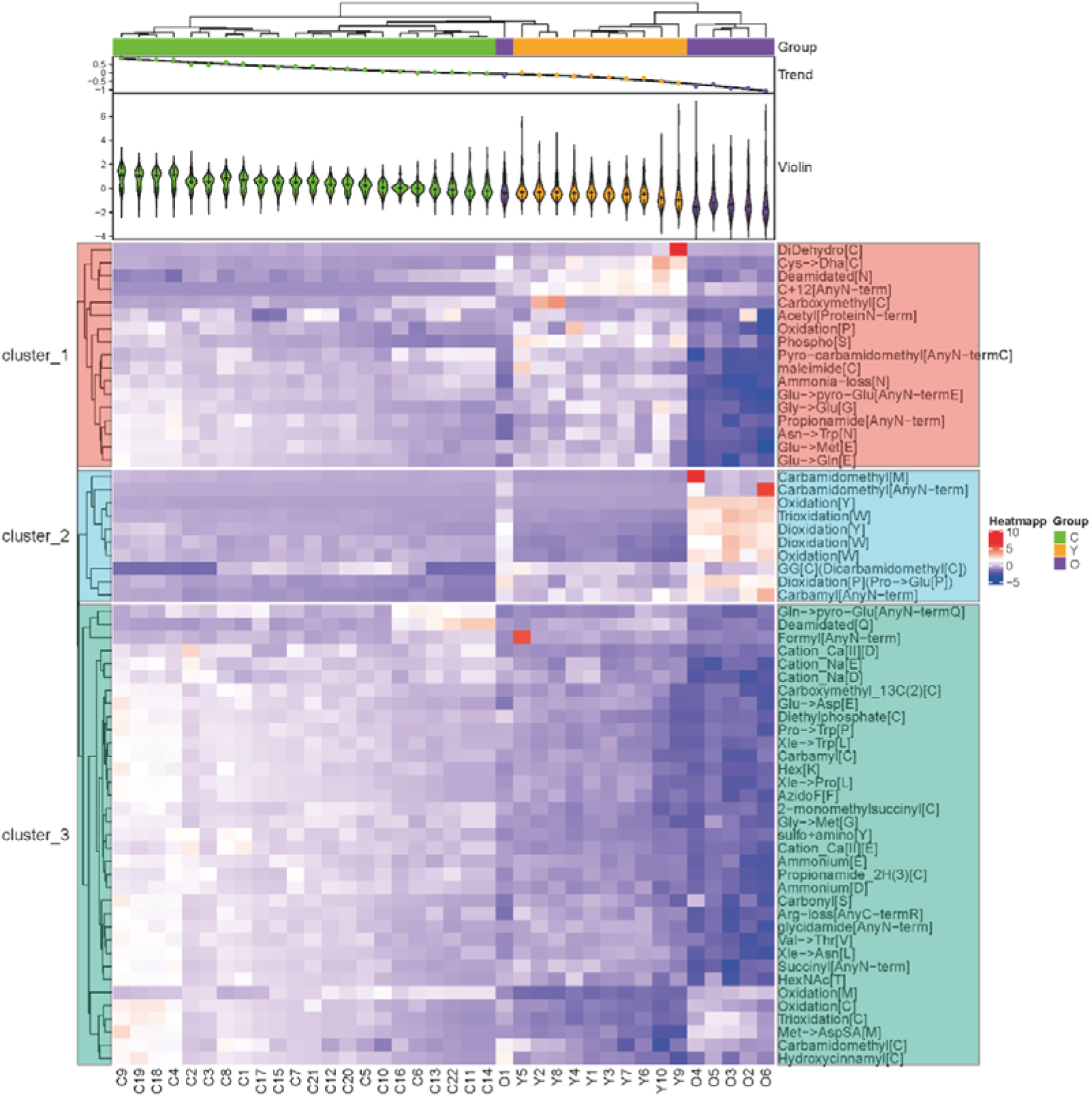
Cluster analysis results of the common modifications among the three groups of samples. Children group covers with green, and young age group covers with orange, the senior group covers with purple.

**Fig S3.**
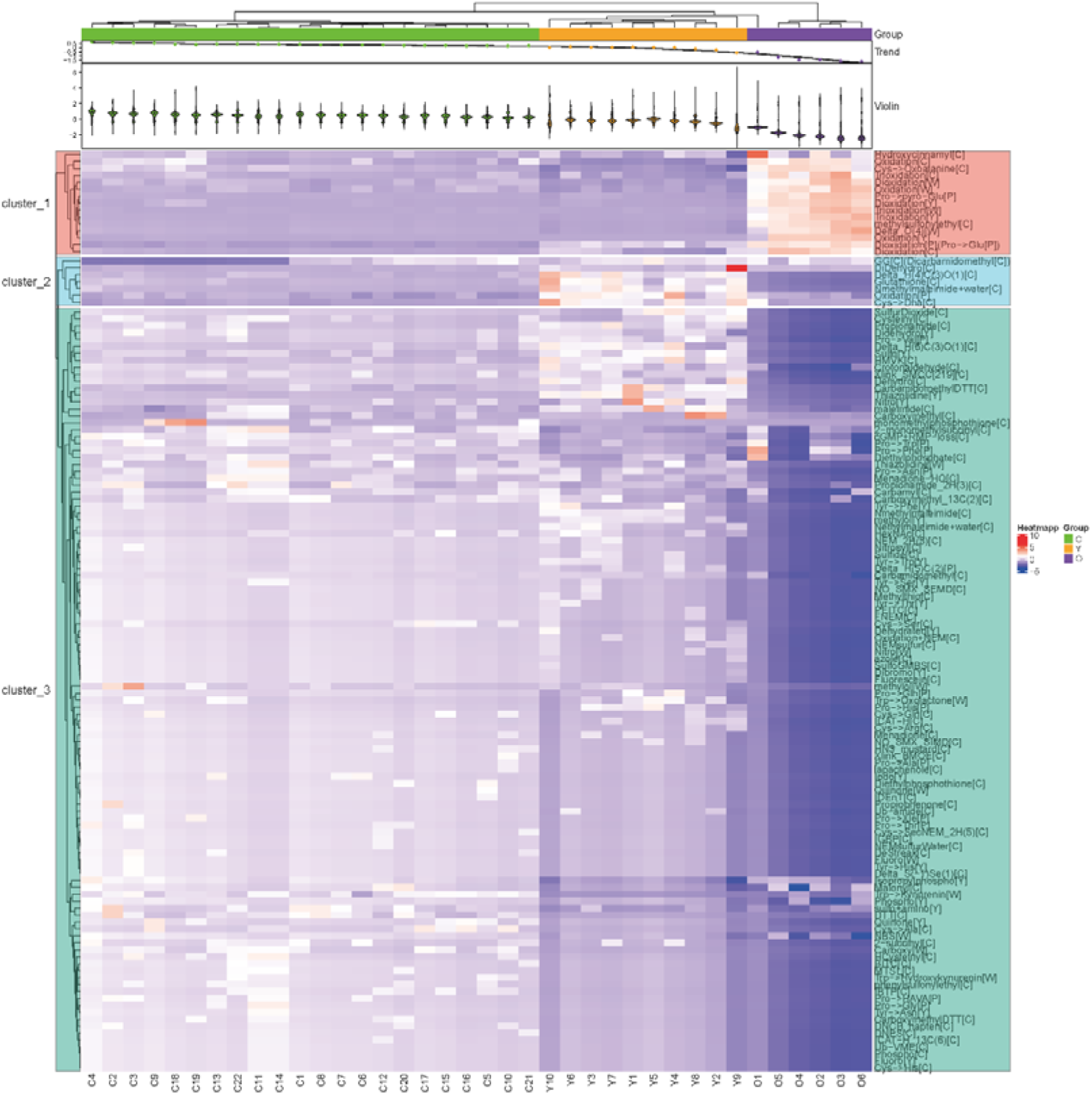
Cluster analysis results of all modifications which occur on proline、tryptophan、tyrosine、cystine among the three groups of samples. Children group covers with green, and young age group covers with orange, the senior group covers with purple.

**Fig S4.**
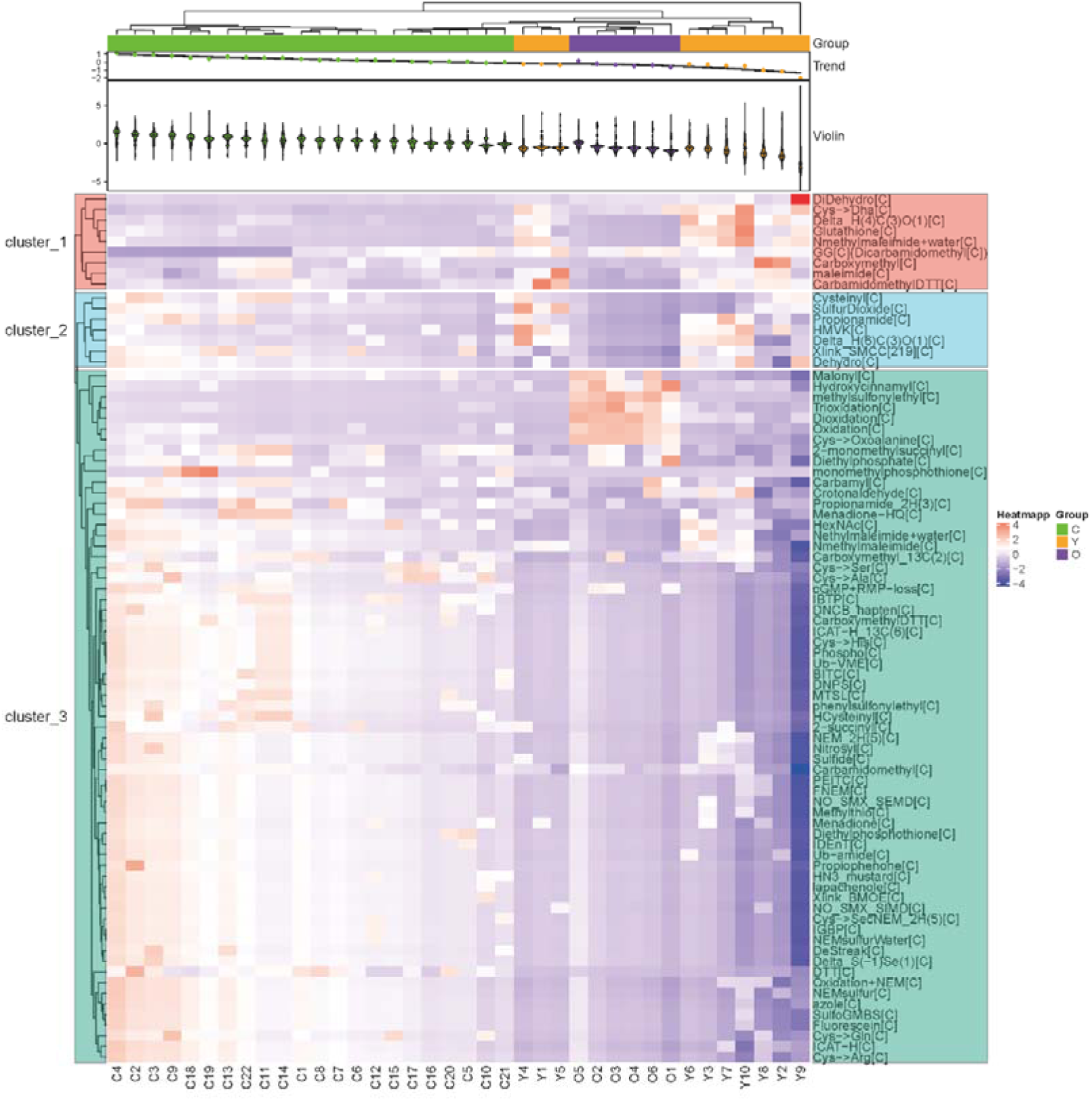
Cluster analysis results of all modifications which occur on cystine among the three groups of samples. Children group covers with green, and young age group covers with orange, the senior group covers with purple.

**Fig S5.**
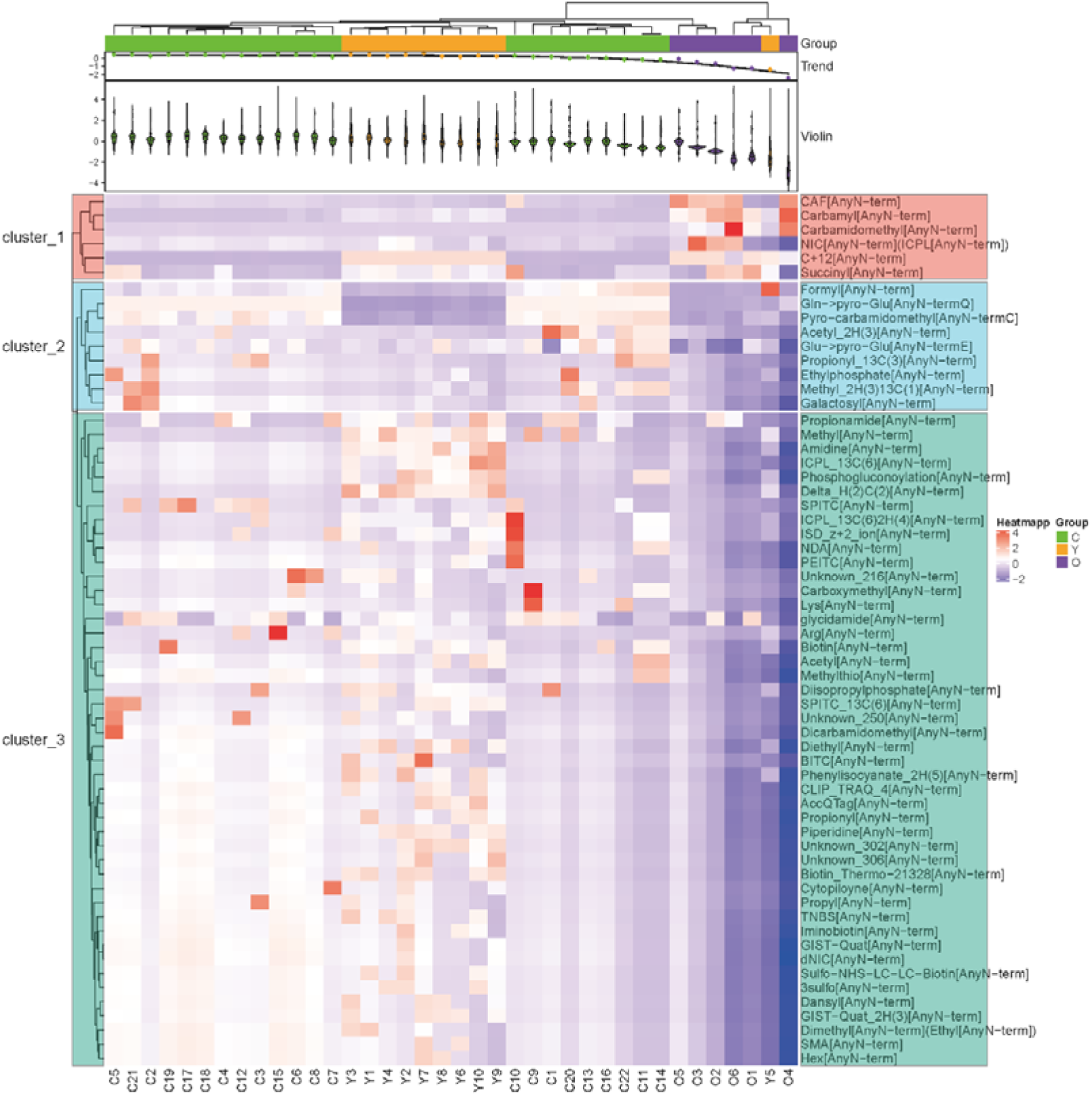
Cluster analysis results of all modifications which occur on anyN-term among the three groups of samples. Children group covers with green, and young age group covers with orange, the senior group covers with purple.

**Supplementary Table 1.**
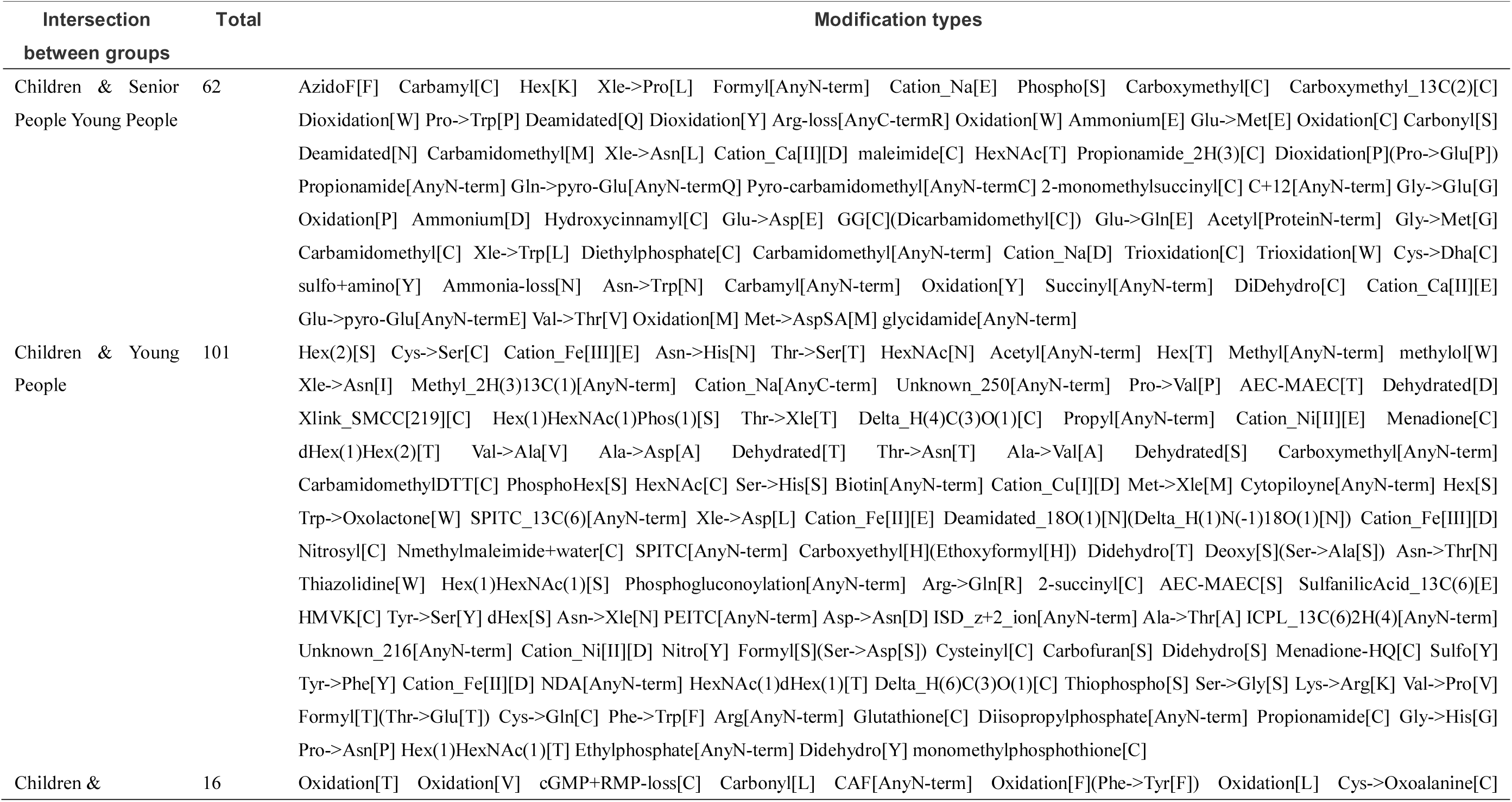
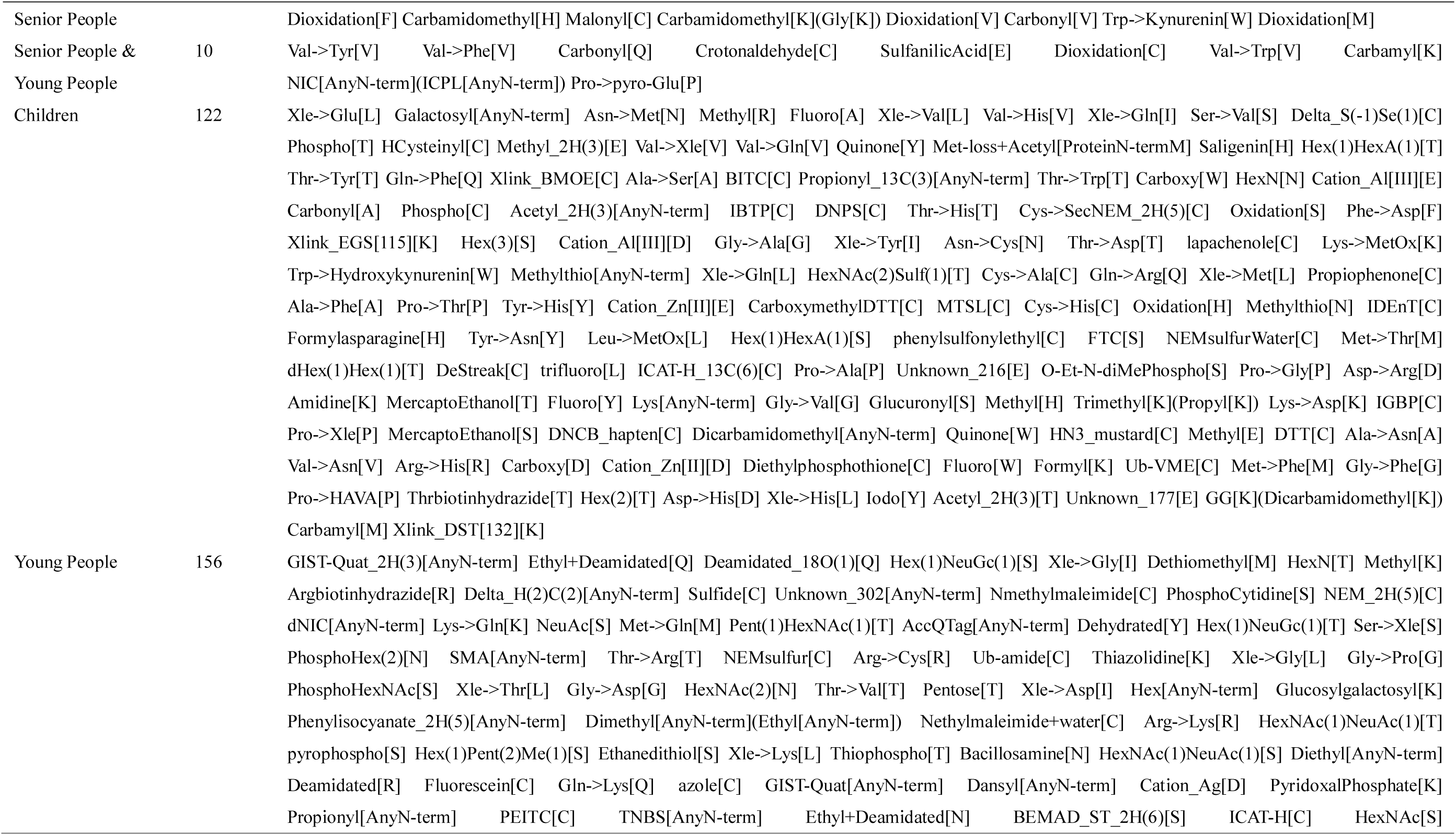
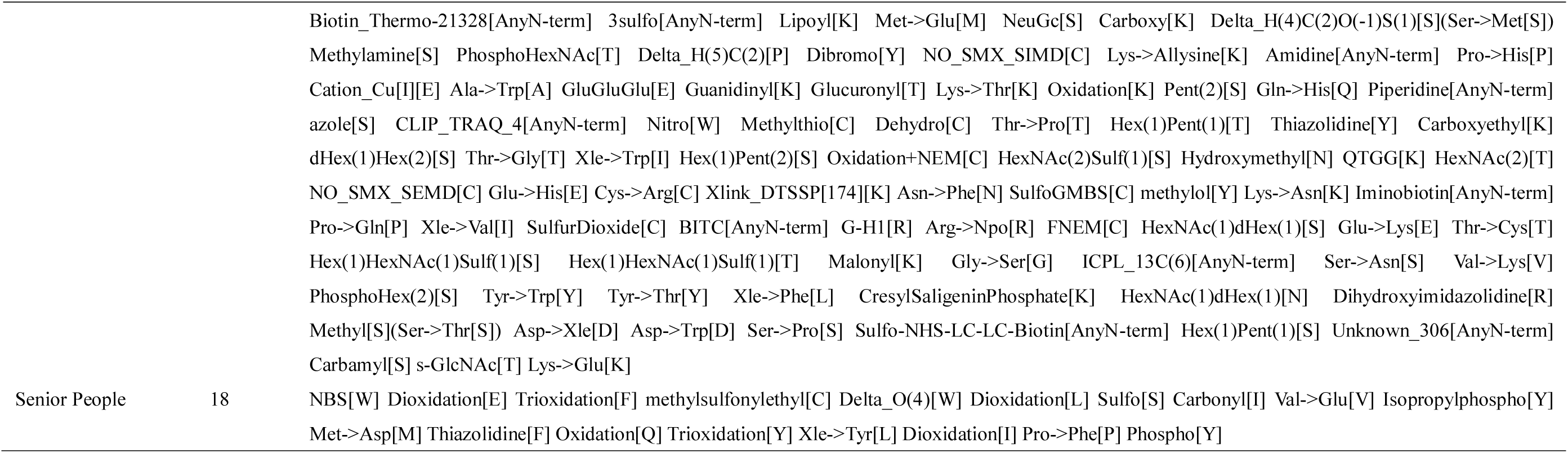
The Venn information of the intersection of sample modification types between different groups

**Supplementary Table 3.**
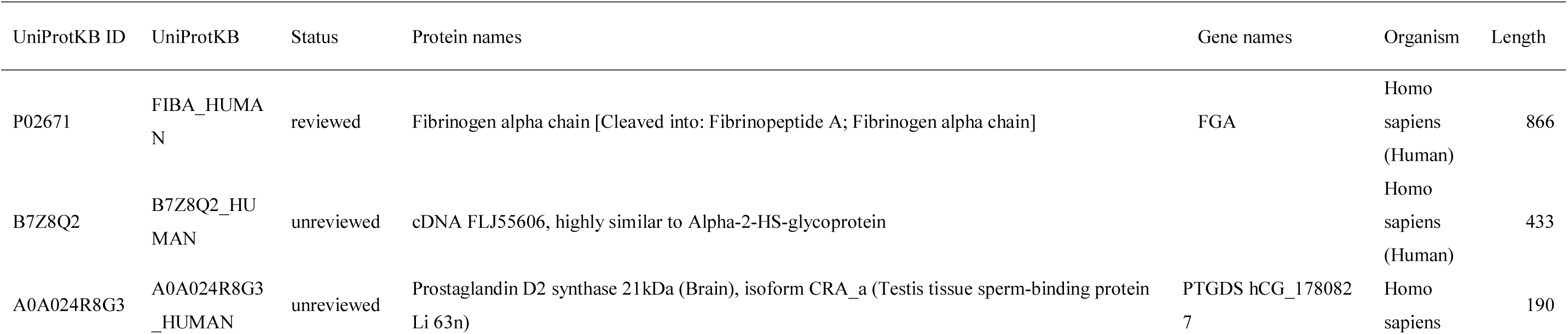
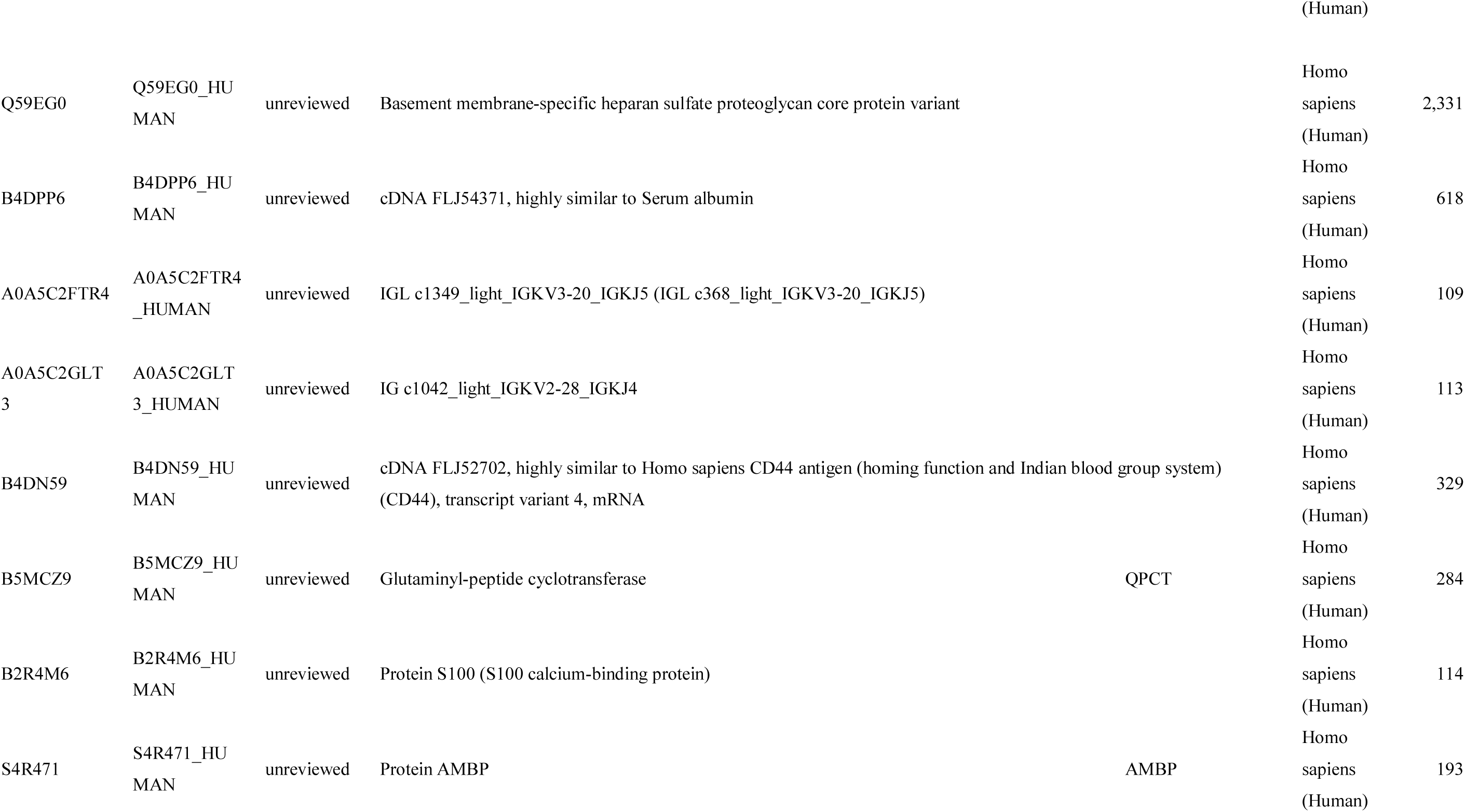
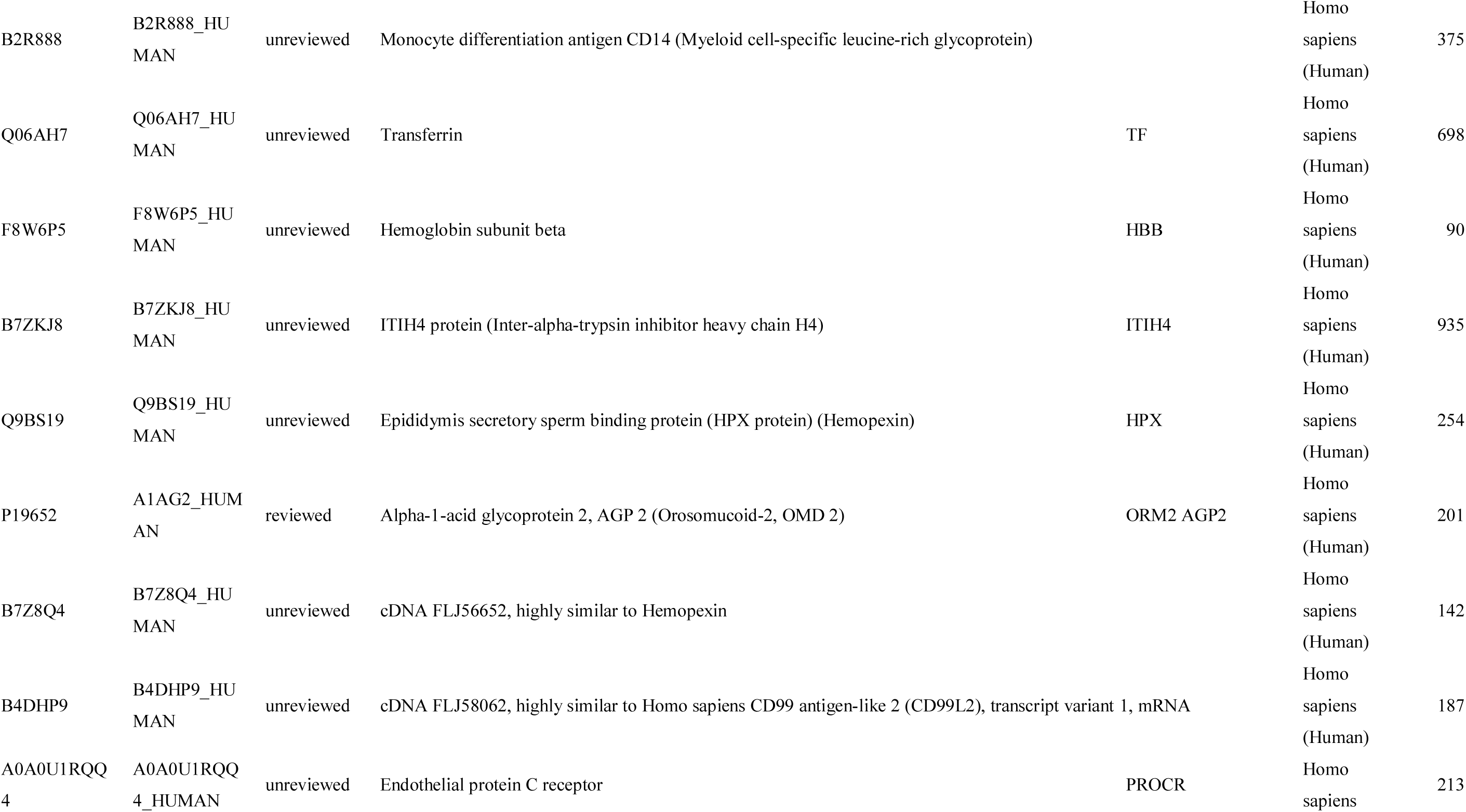
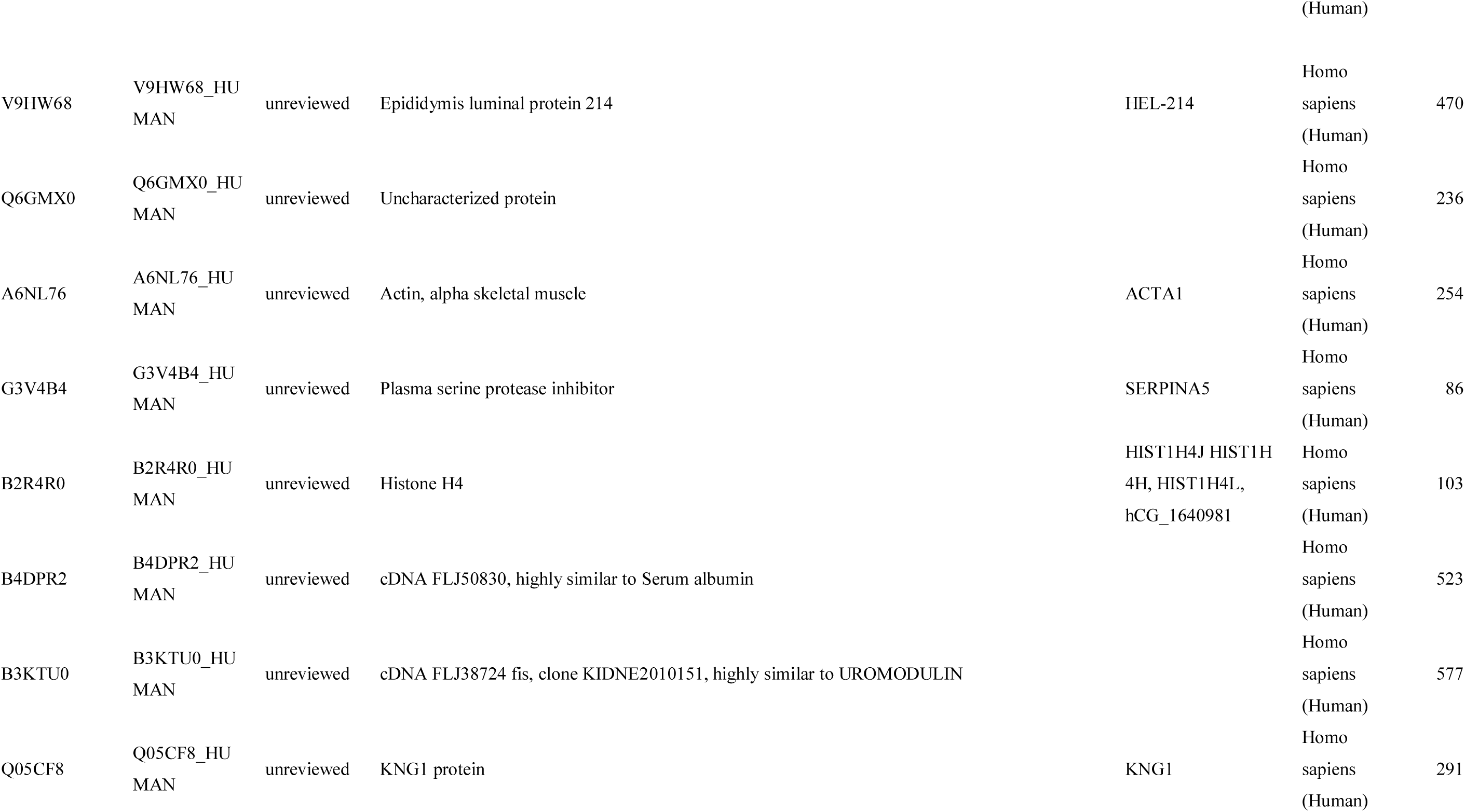
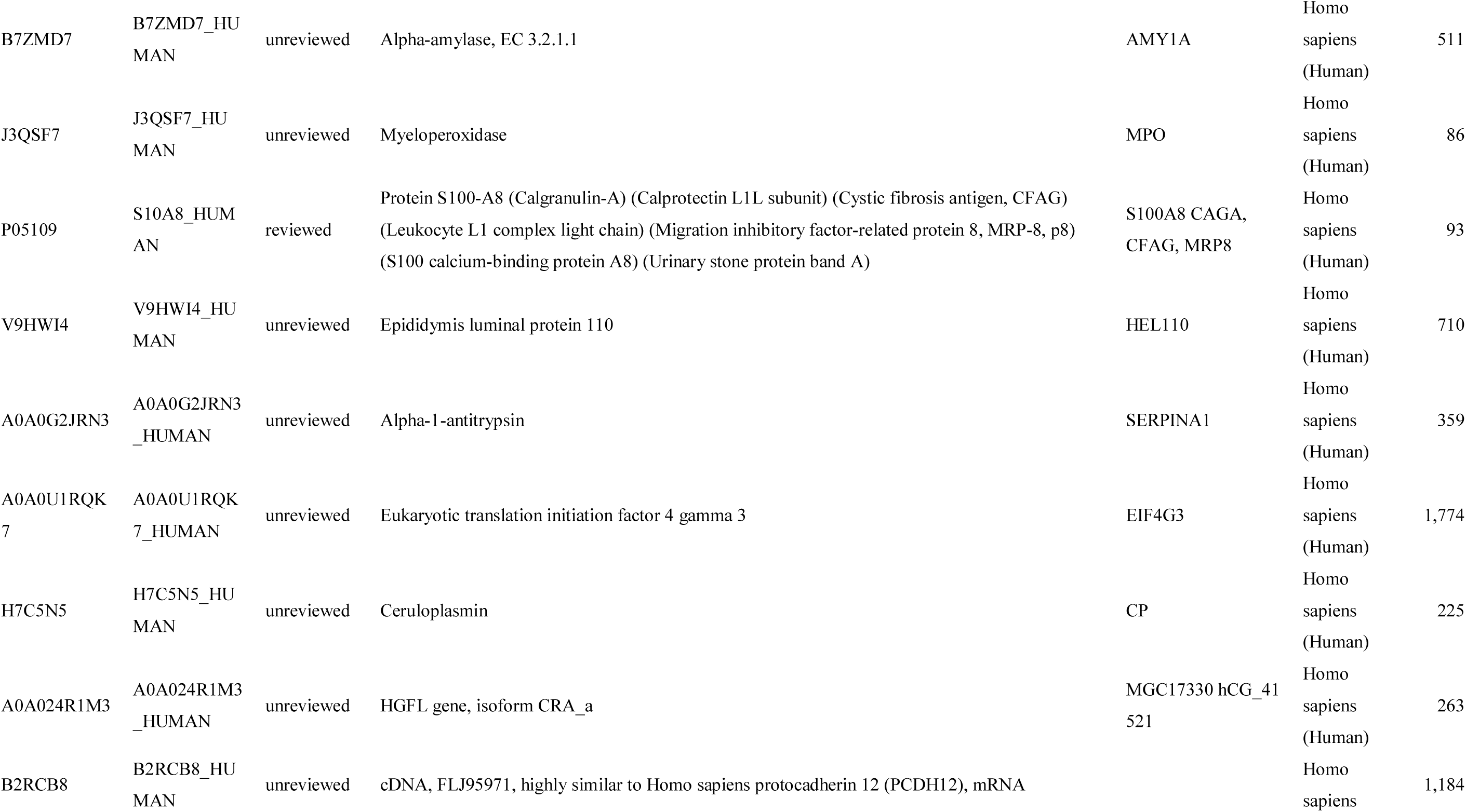
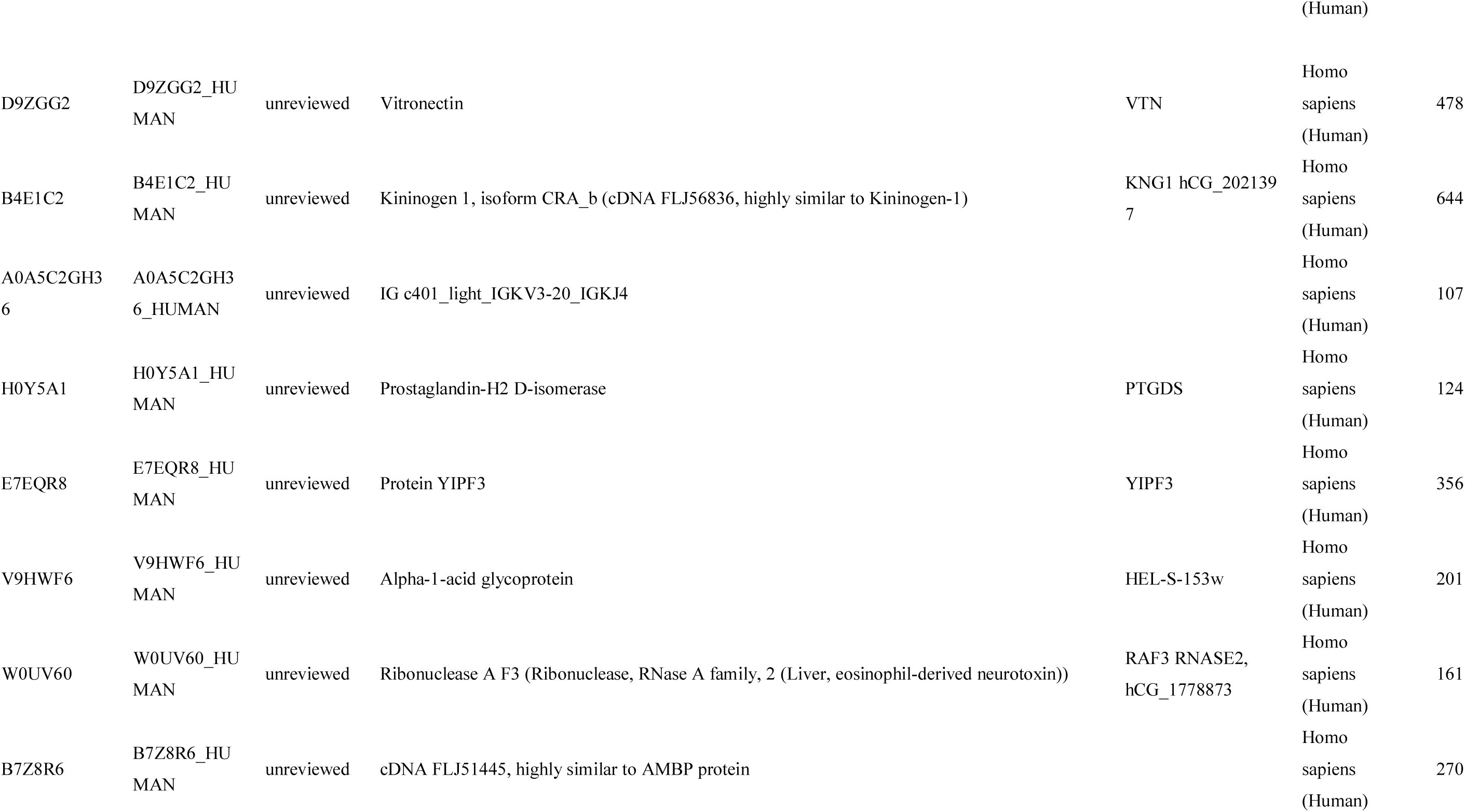
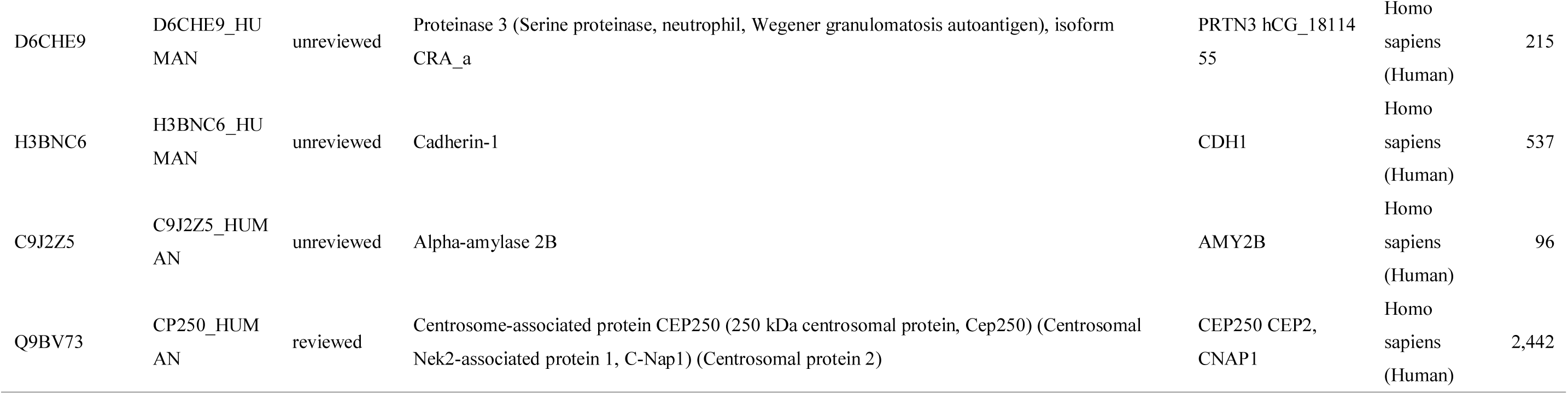
The specific proteins and related information involved in the unique modification of the senior group

